# Fascin limits Myosin activity within *Drosophila* border cells to control substrate stiffness and promote migration

**DOI:** 10.1101/2021.04.27.441651

**Authors:** Maureen C. Lamb, Chathuri P. Kaluarachchi, Thiranjeewa I. Lansakara, Yiling Lan, Alexei V. Tivanski, Tina L. Tootle

**Author notes:** Author Contributions: Conceptualization: M.C.L and T.L.T; Methodology: M.C.L, T.I.L, and A.V.T; Investigation: M.C.L., C.P.K., and T.I.L; Formal Analysis: M.C.L, C.P.K, T.I.L, and Y.L. Writing – Original Draft: M.C.L; Writing – Review and Editing: M.C.L, A.V.T, and T.L.T; Visualization: M.C.L; Funding Acquisition: A.V.T. and T.L.T; Supervision: A.V.T and T.L.T.

## Abstract

A key regulator of collective cell migrations, which drive development and cancer metastasis, is substrate stiffness. Increased substrate stiffness promotes migration and is controlled by Myosin. Using *Drosophila* border cell migration as a model of collective cell migration, we identify, for the first time, that the actin bundling protein Fascin limits Myosin activity *in vivo*. Loss of Fascin results in: increased activated Myosin on the border cells and their substrate, the nurse cells; decreased border cell Myosin dynamics; and increased nurse cell stiffness as measured by atomic force microscopy. Reducing Myosin restores on-time border cell migration in *fascin* mutant follicles. Further, Fascin’s actin bundling activity is required to limit Myosin activation. Surprisingly, we find that Fascin regulates Myosin activity in the border cells to control nurse cell stiffness to promote migration. Thus, these data shift the paradigm from a substrate stiffness-centric model of regulating migration, to uncover that collectively migrating cells play a critical role in controlling the mechanical properties of their substrate in order to promote their own migration. This new means of mechanical regulation of migration is likely conserved across contexts and organisms, as Fascin and Myosin are common regulators of cell migration.

## Introduction

Cell migration is an essential process driving both development and cancer metastasis. During these processes, cells often migrate as groups or collectives, rather than single cells (Friedl and Gilmour, 2009). Collective cell migration requires that cell-cell adhesions be maintained amongst the cells to support cluster cohesion (De Pascalis and Etienne-Manneville, 2017). Additionally, many collective cell migrations occur in an invasive manner with the group of cells migrating between other cells or through basement membranes (Chang et al., 2019). During invasive migration, the environment puts mechanical forces on the migrating cells, causing them to respond by changing their shape and stiffness, and by modifying properties of their environment, such as extracellular matrix (ECM) composition (Aguilar-Cuenca et al., 2014; Eble and Niland, 2019; Gasparski et al., 2017). Therefore stiffness has emerged as a critical regulator of collective cell migration.

During invasive, collective cell migration the group or cluster of cells must generate force necessary to invade through the ECM or other cells. Stiffness of the substrate is considered the primary regulator of the migrating cell’s stiffness and ability to migrate (Aguilar-Cuenca et al., 2014). For example, increased substrate stiffness contributes to cancer cell migration and metastasis (Eble and Niland, 2019; Gasparski et al., 2017; Oakes, 2018). Indeed, hard matrices induce migration in breast cancer cells (Ren et al., 2021), and increased substrate stiffness promotes epithelial to mesenchymal transitions (Nieto and Cano, 2012). While the role of substrate stiffness in promoting cell migration is well-established, most of these studies utilized *in vitro* culture systems. Therefore, it remains poorly understood how migrating cells are regulated in their native environments by the stiffness of their endogenous substrates.

A master regulator of cellular stiffness is Non-Muscle Myosin II (subsequently referred to as Myosin). Myosin is a force generating actin motor (Aguilar-Cuenca et al., 2014; Vicente-Manzanares et al., 2009). It is composed of two copies of three subunits: two heavy chains, two essential light chains, and two regulatory light chains (MRLC; (Aguilar-Cuenca et al., 2014; Vicente-Manzanares et al., 2009)). Myosin activation is regulated through phosphorylation of its regulatory light chains. This phosphorylation occurs through a number of kinases, including Myosin light chain kinase (MLCK) and Rho-associated kinase (Rok), and dephosphorylation occurs through phosphatases, such as protein phosphatase 1c (PP1c) and its catalytic subunit, Myosin binding subunit (Mbs (Aguilar-Cuenca et al., 2014; Vicente-Manzanares et al., 2009)). Myosin generates cortical tension by associating with and acting upon cortical F-actin; this regulates cell stiffness which can influence cell migration (Aguilar-Cuenca et al., 2014; Butcher et al., 2009). Importantly, Myosin regulates stiffness in both substrates and migrating cells during many different cell migrations (Lo et al., 2000; Mohan et al., 2015; Vicente-Manzanares et al., 2009). Additionally, Myosin not only generates mechanical force within a cell but aids in sensing and responding to external forces applied to the cell (Aguilar-Cuenca et al., 2014; Butcher et al., 2009; Vicente-Manzanares et al., 2009).

A recently discovered regulator of Myosin is Fascin. Fascin is an actin binding protein that bundles or cross-links actin filaments into fibers (Hashimoto et al., 2011; Jayo and Parsons, 2010). However, recent studies demonstrate that there are many non-canonical roles for Fascin in the cell (Lamb and Tootle, 2020). One of these non-canonical functions of Fascin is the regulation of Myosin (Elkhatib et al., 2014). Increasing concentrations of Fascin in an *in vitro* system decreased Myosin ATP consumption and motor speed along actin filaments (Elkhatib et al., 2014). These data suggest that Fascin limits Myosin activity (Elkhatib et al., 2014). Whether Fascin limits Myosin activity to control substrate stiffness and thereby cell migration remains unknown. Notably, Fascin has well-established roles in promoting cell migration (Lamb and Tootle, 2020). Fascin aids in the formation of cell migratory structures like filopodia (Hashimoto et al., 2011) and invadopodia (Li et al., 2010). Fascin promotes many types of cell migrations in development and disease, including cancer metastasis (Ma and Machesky, 2015). Investigation of Fascin’s role in promoting cell migration has primarily focused on Fascin as an actin bundler and it is unknown if Fascin limits Myosin activity to regulate collective cell migration.

An ideal model to uncover the role of Fascin in regulating Myosin during collective cell migration in a native context is *Drosophila* border cell migration. Border cell migration occurs during Stage 9 (S9) of oogenesis. During S9, the follicle is composed of an oocyte and 15 germline-derived nurse cells that are surrounded by a layer of somatic epithelial cells called follicle cells (Spradling, 1993). At the beginning of S9, a group of 8-10 outer follicle cells are specified as border cells and delaminate from the epithelium to start their migration (Montell, 2003). The border cells migrate invasively and collectively between the nurse cells – which are the substrate for the migration – until they reach the nurse cell-oocyte boundary (Montell, 2003). Importantly, similar to other types of migration, the stiffness of the nurse cell substrate regulates both the stiffness of the border cells and their migration (Aranjuez et al., 2016). Therefore, border cell migration is a powerful model for studying invasive, collective cell migration as the cluster of cells can be visualized in its native context using both fixed and live imaging. Additionally, the factors that regulate border cell migration play conserved roles in other invasive, collective cell migrations, including cancer metastasis (Montell et al., 2012; Stuelten et al., 2018). Indeed, both Fascin and Myosin play roles in promoting cancer metastasis (Aguilar-Cuenca et al., 2014; Hashimoto et al., 2011) and on-time border cell migration (Edwards and Kiehart, 1996; Lamb et al., 2020). We previously found that Fascin (*Drosophila* Singed, Sn) is required for both border cell delamination and proper protrusion localization (Lamb et al., 2020). Both loss and activation of Myosin result in similar phenotypes (Aranjuez et al., 2016; Majumder et al., 2012; Mishra et al., 2019). These data suggest that the cycling of Myosin between active and inactive forms controls border cell migration. Thus, border cell migration is an ideal system to uncover the relationship of Fascin and Myosin during collective cell migration.

Here, we demonstrate for the first time that Fascin inhibits Myosin activity *in vivo*. Loss of Fascin significantly increases the level of active Myosin, reduces Myosin dynamics, and increases nurse cell (aka substrate) stiffness as quantified by atomic force microscopy (AFM) nanoindentation technique. Reducing Myosin in *fascin* mutant follicles rescues border cell migration delays, indicating that Fascin’s tight regulation of Myosin activity is critical for on-time migration. Further, a phosphomimetic form of Fascin that precludes bundling is unable to limit Myosin activation, supporting the prior model that Fascin limits Myosin activity by tightly bundling actin and preventing Myosin binding to actin filaments. We used RNAi knockdown and rescue experiments to assess the cell-specific roles of Fascin in regulating Myosin activity and nurse cell stiffness. Based on the literature, we expected that Fascin would primarily function within the nurse cells to control both substrate stiffness and myosin activity within both the nurse cells and border cells. Surprisingly, we find that knocking down Fascin in the border cells increases the level of active Myosin on both the border cells and the nurse cells, and increases the stiffness of the nurse cells. Similarly, re-expressing Fascin in only the border cells of *fascin* mutants restores normal Myosin activity levels and stiffness of the nurse cells. These unexpected and novel findings suggest that migrating cells influence the mechanobiology of their substrate to promote their migration. Supporting this, increasing Rok activity in the border cells also results in increased nurse cell stiffness, indicating this migratory cell regulation of substrate stiffness is not a Fascin specific phenomenom. Together these findings lead to the following model: Fascin acts primarily within the migrating border cells to limit Myosin activation which controls the stiffness of the both the border cells and their substrate, the nurse cells, to promote on-time migration. It is likely that this novel regulation of Myosin by Fascin and thereby, the migrating cells controlling substrate stiffness is a conserved means of promoting other collective cell migrations.

## Results

### Fascin inhibits Myosin activation in the *Drosophila* follicle

Previous data demonstrates that Fascin can inhibit the activity of Myosin *in vitro* (Elkhatib et al., 2014). Fascin functions in both the nurse cells and the border cells to promote on-time border cell migration (Lamb et al., 2020). Additionally, during border cell migration Myosin generates forces in the nurse cells that push upon the border cells, causing them to activate Myosin and stiffen (Aranjuez et al., 2016), suggesting that the nurse cells control the stiffness of the border cell cluster. Based on these observations, we hypothesized that Fascin may regulate Myosin activity in the *Drosophila* follicle, specifically the nurse cells, to promote border cell migration.

To test this hypothesis, we assessed if Fascin limits Myosin activity in the *Drosophila* follicle. Myosin is activated via phosphorylation on its regulatory light chain subunit (MRLC). To assess changes in Myosin activation in the follicle, we stained follicles using an antibody against phosphorylated MRLC (pMRLC); wild-type and *fascin*-null follicles were stained in the same tube to account for staining variability. We observe a striking increase in active MRLC along both the nurse cell and border cell membranes of *fascin*-null follicles (Figure 1B compared to A, orange arrows and B’ compared to A’, blue arrows). We quantified levels of active MRLC by measuring the relative fluorescent intensity of pMRLC on the nurse cell and border cell membranes (Figure 1C-D, see Methods for quantification details); in all graphs of MRLC activity nurse cell measurements are shown in orange, while border cell measurements are shown in blue. There is a significant increase in active MRLC intensity on the *fascin*-null nurse cell membranes compared to wild-type follicles (Figure 1C, p<0.0001). Additionally, active MRLC is also significantly increased on the border cell cluster when Fascin is lost (Figure 1D, p<0.0001). We also quantified changes in active MRLC puncta number and length on the border cells cluster (Figure 1 E, F; see Methods for quantification details). Loss of Fascin increases puncta number but decreases puncta length (Figure 1 E, F, p<0.001). Together these results demonstrate that Fascin limits Myosin activation in the *Drosophila* S9 follicle on both the nurse cell membranes and the border cell cluster, providing the first evidence that Fascin regulates Myosin activity *in vivo*.

**Figure 1:**
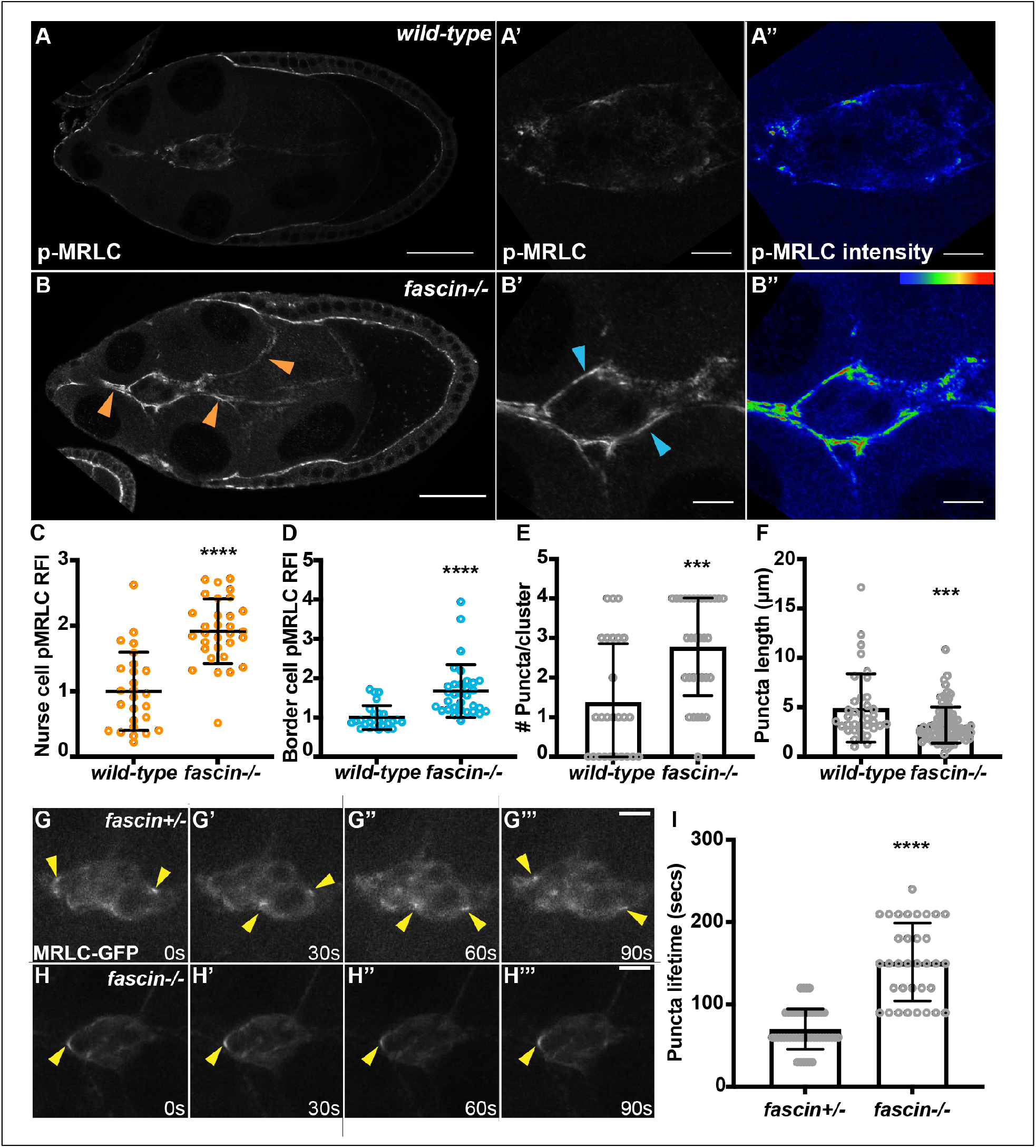
Fascin limits myosin activity in the Stage 9 *Drosophila* follicle. (**A-B’’**) Maximum projections of 2-4 confocal slices of Stage 9 follicles of the indicated genotypes. (**A-A’, B-B’**) phospho-MRLC (pMRLC, white). (**A’’, B’’**) phospho-MRLC (pMRLC) pseudocolored with Rainbow RGB, red=highest intensity pixels. (**A-A’’**) wild-type (*yw*). (**B-B’’**) *fascin*-null (*fascin*^*sn28/sn28*^). Samples were stained in the same tube. Orange arrows = pMRLC enrichment on nurse cells. Blue arrows = pMRLC enrichment on border cell cluster. Scale bars = 50μm in **A, B** and 10μm in **A’-A’’, B’-B’’**. (**C-F**) Graphs of quantification of pMRLC intensity and localization at the nurse cell membranes (**C**) and border cell cluster (**D, E, F**) in wild-type and *fascin*-null follicles. Each circle represents a follicle. Error bars=SD. ***p<0.001, ****p<0.0001 (unpaired t-test). In **C**, peak pMRLC intensity was quantified at the nurse cell membranes and normalized to phalloidin staining in the same follicle, three measurements were taken per follicle and averaged. In **D**, pMRLC intensity on the border cell cluster was quantified and normalized to background staining in the same follicle. In **E**, the number of myosin puncta per cluster was manually counted. In **F**, the length of each myosin puncta was measured. (**G-H’’’**) Maximum projection of 3 confocal slices from time-lapse imaging of MRLC-GFP expression in the indicated genotypes. Direction of migration is to the right. Scale bars= 10μm. (**G-G’’’**) Control follicle (*fascin*^*sn28*^*/+*; *GFP-MRLC/+*; Video 1). (**H-H’’’**) *fascin*-null follicle (*fascin*^*sn28/sn28*^; *GFP-MRLC/+*; Video 2). (**I**) Quantification of puncta lifetime from time-lapse imaging for control (n=4) and *fascin*-null (n=4) GFP-MRLC expressing follicles. Puncta lifetime was defined as the amount of time elapsed from when a punctum first appeared to when it completely disappeared. ****p<0.0001 (unpaired t-test). Error bars=SD. *fascin*-null follicles have increased pMRLC on the the nurse cell membranes (B, C) and border cell cluster (B’, D) compared to wild-type follicles (A, A’, C, D). The border cell clusters in *fascin*-null mutants also have increased Myosin puncta number but decreased length (E, F). *fascin* mutants have significantly slowed Myosin dynamics (H-H’’’, I) compared to the control clusters (G-G’’’, I).

### Fascin limits Myosin dynamics on the migrating border cell cluster

We next wanted to determine how Fascin influences Myosin dynamics during border cell migration. In addition to the level of activation, the localization and dynamics of Myosin influence invasive migration (Aguilar-Cuenca et al., 2014; Aranjuez et al., 2016; Majumder et al., 2012; Vicente-Manzanares et al., 2009). Indeed, during border cell migration, dynamic cycles of Myosin activation and inactivation at the cluster membrane are essential for proper migration (Aranjuez et al., 2016). We visualized Myosin dynamics on the border cell cluster using a C-terminally GFP-tagged MRLC (*Drosophila* Spaghetti Squash, Sqh), under the control of its endogenous promoter. Previous data demonstrates that MRLC-GFP is highly expressed on the border cell cluster during migration and accumulates in transient puncta on the cluster; these puncta depend on Myosin activation, suggesting they are sites of active Myosin (Majumder et al., 2012). Using live imaging, we find in control follicles, MRLC-GFP puncta appear and disappear rapidly on the border cell cluster (Figure 1G-G’’’, Video 1). However, in the *fascin*-null follicles, the MRLC-GFP puncta dynamics are much slower (Figure 1H-H’’’, Video 2). We quantified this change in MRLC-GFP dynamics by measuring puncta lifetime on the cluster (Figure 1I). The control follicles display an average puncta lifetime of 70.2 seconds, while in *fascin*-null follicles the average puncta lifetime is 151.8 seconds (Figure 1I, p<0.0001). These results suggest that Fascin limits Myosin dynamics on the migrating border cell cluster.

### Fascin regulates nurse cell stiffness

As increased Myosin activity increases actomyosin contractility and cell stiffness, we next wanted to directly measure the stiffness of *fascin*-null follicles. Substrate stiffness is thought to be a driving regulator of cell migration and migrating cell stiffness (Di Martino et al., 2016; Gasparski et al., 2017; Oakes, 2018; Ren et al., 2021), therefore we aimed to directly quantify nurse cell stiffness. AFM is a standard method to directly measure mechanical properties of biological tissues (Kreplak, 2016). AFM can be used to quantify the elastic modulus, which is a measurement of how easily an elastic material is deformed when a known amount of force is applied (Kreplak, 2016). A high elastic modulus value corresponds to a stiff tissue. We used AFM nanoindentation technique to quantify the stiffness of *fascin*-null and wild-type nurse cells (Chen et al., 2019; Crest et al., 2017). The nurse cells are the substrate for the border cells and their stiffness regulates border cell migration and cluster stiffness (Aranjuez et al., 2016). Notably, during S9, the nurse cells are surrounded by a layer of stretch follicle cells and a basement membrane that envelopes the entire follicle (Figure 2A). Previous measurements on *Drosophila* follicles using AFM established that there is significant difference in stiffness between the basement membrane and the underlying nurse cells (Chen et al., 2019; Crest et al., 2017). These different tissues stiffnesses can be separated by using different indentation ranges to indent the AFM probe into just the basement membrane or to indent deeper into the nurse cells (Figure 2B; (Chlasta et al., 2017)). Thus, by using two indentation ranges to fit the mechanical response we can quantify the distinct stiffness of the basement membrane versus that of the underlying nurse cells (Chlasta et al., 2017).

**Figure 2:**
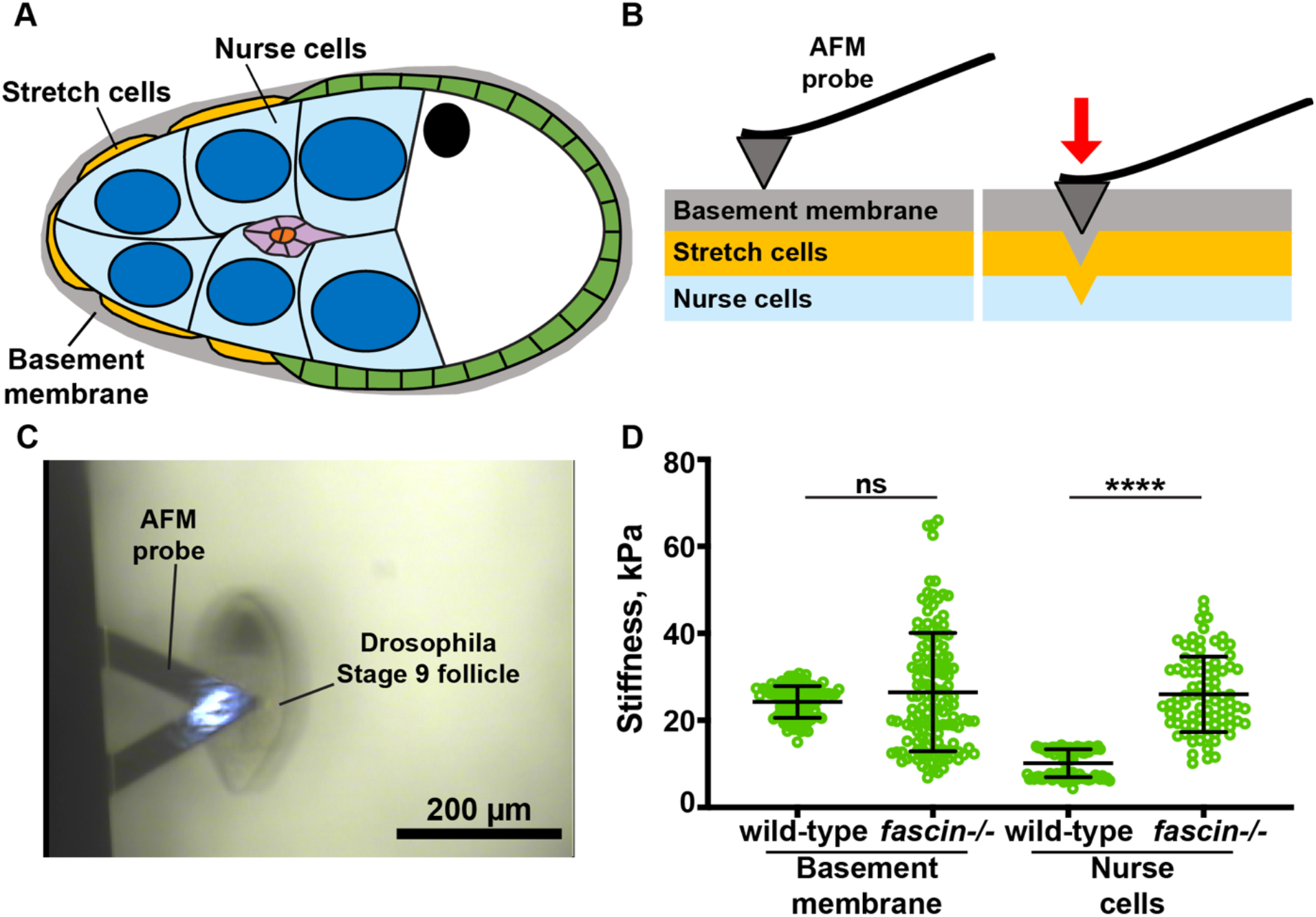
Fascin regulates nurse cell stiffness in the *Drosophila* follicle. (**A**) Schematic of Stage 9 Drosophila follicle. The nurse cells (light blue) are surrounded by a layer of stretch cells (gold) and basement membrane (grey). (**B**) Schematic of AFM probe indentation through the basement membrane (grey) and stretch cells (gold) into the underlying nurse cells (light blue). (**C**) Bright field image of AFM probe over a S9 follicle. (**D**) Graph of nurse cell stiffness (kPa) in wild-type or *fascin*-null follicles as measured by AFM. Each circle represents a single indentation. Error bars=SD. ns indicates p>0.05, ****p<0.0001 (unpaired t-test). Loss of Fascin significantly increases the stiffness of the nurse cells (D).

We use AFM and the Hertzian elastic contact model to calculate the stiffness of wild-type and *fascin*-null S9 follicles (Figure 2C); for increased clarity, graphs quantifying stiffness are represented in green. For an indentation range of 20-200nm, which probes the basement membrane, wild-type follicles have an average stiffness of 24.2 kPa and *fascin*-null follicles have a similar average stiffness of 26.5 kPa (Figure 2D, p>0.05). However, for an indentation range of 200-800nm, which probes the nurse cell stiffness, wild-type follicles have an average stiffness of 10.1 kPa while *fascin*-null follicles have a significantly increased average stiffness of 25.9 kPa (Figure 2D, p<0.0001). Thus, the stiffness of the *fascin*-null nurse cells is >2x higher than wild-type nurse cells. Together these results demonstrate that loss of Fascin increases the stiffness of the nurse cells in S9 *Drosophila* follicles.

### Fascin limits Myosin activity to promote border cell migration

As increased stiffness of the nurse cells or border cells leads to border cell migration delays (Aranjuez et al., 2016), we hypothesized that the increased Myosin in *fascin*-null follicles contributes to the previously characterized border cell migration delays (Lamb et al., 2020). To address this hypothesis, we first used pharmacological inhibitors of Myosin and assessed their effects on border cell migration. Follicles were incubated for 2 hours in either control media or 200µM of Y-27632, a Rho inhibitor previously used to reduce Myosin activity in *Drosophila* follicles (He et al., 2010), or 200µM of blebbistatin, a Myosin inhibitor. These inhibitors reduce activated Myosin levels on both the nurse cells and border cells (Figure 3-figure supplement 1A, B). We then employed our previously developed method to quantify delays in border cell migration during S9, which takes the ratio of the distance the border cells have migrated from the anterior end of the follicle to the distance of the outer follicle cells from the anterior end of the follicle (see schematic Figure 3G (Lamb et al., 2020)). We call this value the migration index; to increase clarity, all migration indexes data are in magenta. A migration index of approximately 1 indicates on-time migration during S9, while a value less that 1 indicates delayed migration and a value greater than 1 indicates an accelerated migration. As we previously established, loss of Fascin significant delays migration (Lamb et al., 2020). Here we find that inhibiting Myosin activity with either drug in *fascin*-null follicles restores on-time border cell migration compared to the *fascin*-null control (Figure 3B-D, H, migration index 1.1 and 1.0 compared to 0.78) and is not significantly different from the wild-type control (Figure 3A, H).

**Figure 3:**
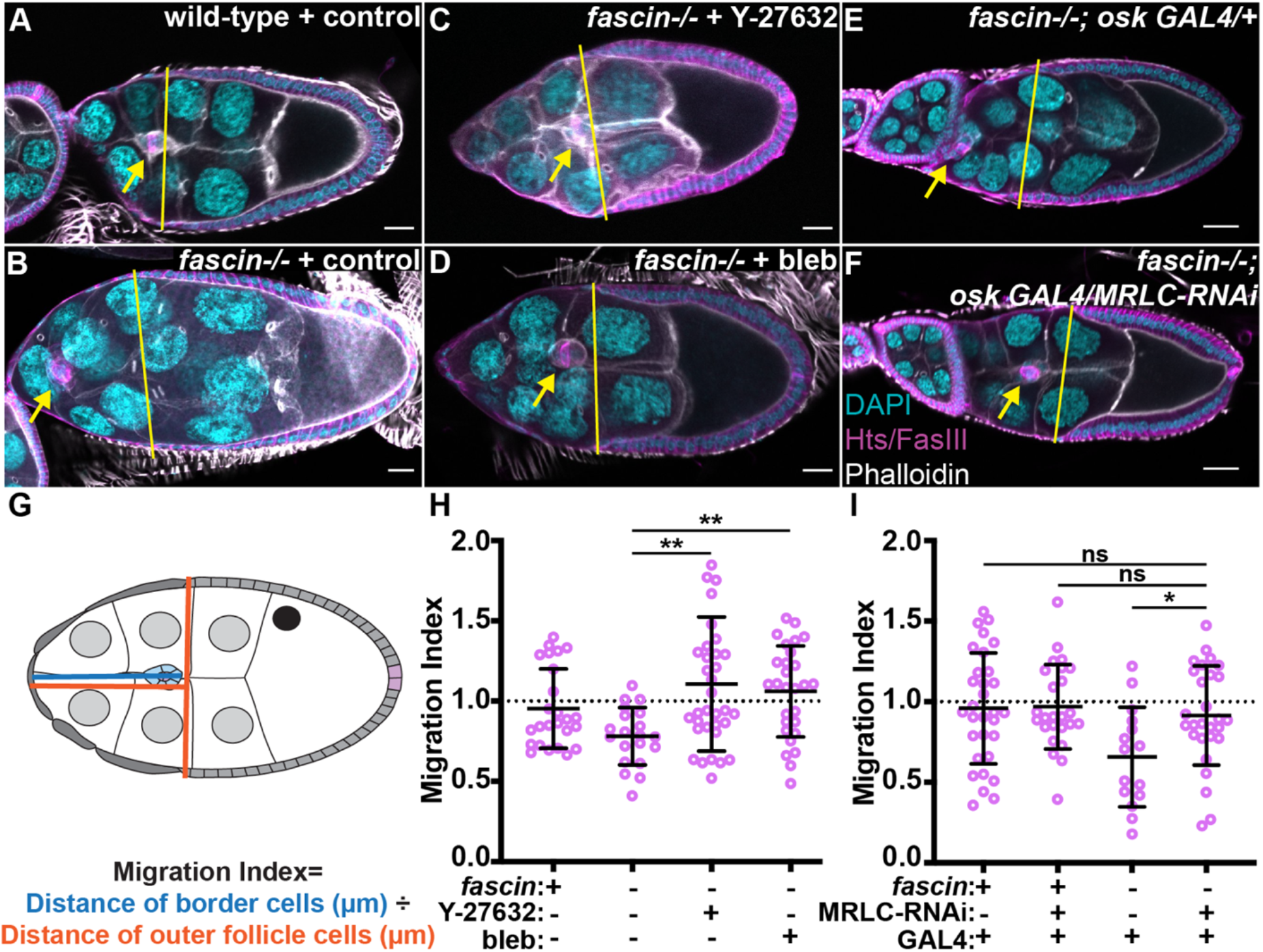
Reducing Myosin activity rescues border cell migration in *fascin* mutant follicles. (**A-F**) Maximum projections of 2-4 confocal slices of Stage 9 follicles of the indicated genotypes. Merged images: Hts/FasIII (magenta, border cell migration stain), phalloidin (white), and DAPI (cyan). Yellow lines = outer follicle cell distance. Yellow arrows = border cell cluster. Black boxes have been added behind text. Scale bars = 20 μm. (**A**) wild-type (*yw*) treated with control S9 media + vehicle (DMSO). (**B**) *fascin-/-*(*fascin*^*sn28/sn28*^) treated with control S9 media. (**C**) *fascin-/-*(*fascin*^*sn28/sn28*^) treated with 200µM of Y-27632. (**D**) *fascin-/-*(*fascin*^*sn28/sn28*^) treated with 200µM of blebbistatin (bleb). (**E**) *fascin*^*sn28/sn28*^; *oskar GAL4 (2)/+* (**F**) *fascin*^*sn28/sn28*^; *oskar GAL4 (2)/MRLC-RNAi*. (**G**) A schematic of the migration index quantification for border cell migration during S9. The migration index is the distance the border cell cluster has migrated divided by the distance of the outer follicle cells from the anterior end. A value of ∼1 indicates on-time migration, a value <1 indicates delayed migration and a value >1 indicates accelerated migration. (**H, I**) Migration index quantification of the indicated genotypes. Dotted line at 1 = on-time migration. Circle = S9 follicle. Lines = averages and error bars = SD. ns indicates p>0.05, *p<0.05, **p < 0.01 (One-way ANOVA with Tukey’s multiple comparison test). Pharmacological inhibition of Myosin rescues border cell migration delays in *fascin* mutant follicles (A-D, H). Similarly, germline knockdown of MRLC restores on-time border cell migration in *fascin* mutants, suggesting increased active Myosin in *fascin* mutants leads to border cell migration delays (E, F, I).

As loss of Fascin increases Myosin activity in both the nurse cells and the border cells, we next sought to identify if this increase in Myosin activity leads to delays in on-time border cell migration. We used the UAS/GAL4 system to express an RNAi against MRLC (*Drosophila* Sqh) to knockdown Myosin in *fascin*-null mutants in different cell types – the germline (*matα* GAL4), somatic (*c355* GAL4), or border cells (*c306* GAL4). Unfortunately, knockdown of Myosin in the somatic (*c355* GAL4) or border cells (*c306* GAL4) was lethal, however knockdown of Myosin in the germline (*matα* GAL4) was viable. Germline knockdown of MRLC in *fascin* mutants significantly decreased active Myosin levels on the nurse cells compared to *fascin*-null controls (Figure 3-figure supplement 1C). However, it fails to restore normal levels of active Myosin on the border cell cluster, as Myosin activation remains significantly increased compared to the wild-type control and is not signficantly different than the *fascin*-null control (Figure 3-figure supplement 1D). We next assessed whether altering Myosin activity within the nurse cells can restore on-time border cell migration in *fascin-*null mutants using the migration index quantification (see schematic, Figure 3G). Germline knockdown of MRLC in *fascin* mutant follicles rescues border cell migration (Figure 3E, F, I, migration index 0.91 compared to 0.65). Together these results suggest that Fascin is required to limit Myosin activity within the nurse cells to promote on-time border cell migration.

### Phosphorylation of Fascin controls its ability to limit Myosin activity

Previous data demonstrated that *in vitro* Fascin can limit Myosin activation, however the mechanism of how Fascin regulates Myosin activity is unknown (Elkhatib et al., 2014). It was hypothesized that Fascin’s ability to tightly bundle actin precludes Myosin from being able to bind to actin filaments and generate force (Elkhatib et al., 2014). Phosphorylation of Fascin at serine 52 (S52, mammalian S39) inhibits its actin bundling function (Ono et al., 1997; Yamakita et al., 1996). If Fascin’s actin bundling activity is required to limit Myosin activation, we would predict that global expression (*actin 5c* GAL4) of phosphomimetic Fascin (S52E) in *fascin*-null mutants would fail to suppress the increased active Myosin. As a control, we find that global expression of wild-type Fascin (GFP-Fascin) significantly reduces active MRLC enrichment on both the nurse cell membranes (Figure 4B, D) and border cell cluster (Figure 4B, blue arrows and E). Conversely, when phosphomimic form of Fascin (GFP-Fascin S52E) is expressed in *fascin*-null mutants we observe high levels of active Myosin on both the nurse cell membranes (Figure 4C, orange arrows and D) and border cell cluster (Figure 4C, blue arrows and E) that are not significantly different than the *fascin* mutant control (Figure 4A, D, E). These data support the model that Fascin limits Myosin activity by bundling actin and precluding Myosin’s ability to bind to actin filaments.

**Figure 4:**
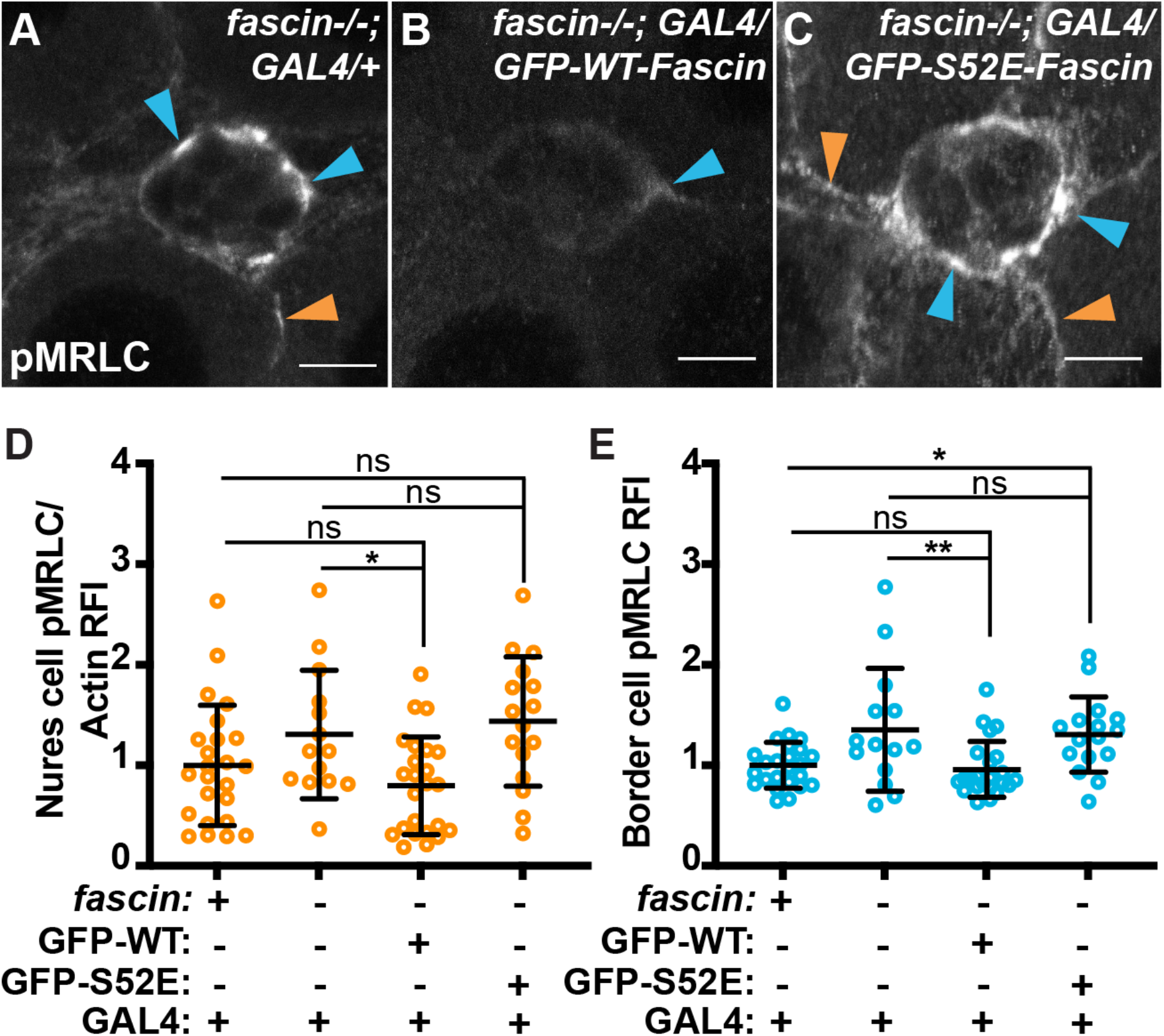
Phosphorylation of Fascin regulates Myosin activation. (**A-C**) Maximum projections of 2-4 confocal slices of Stage 9 follicles of the indicated genotypes stained for phospho-MRLC (pMRLC, white). Orange arrows = pMRLC enrichment on surrounding nurse cells. Blue arrows = pMRLC enrichment on border cell cluster. Scale bars = 10μm. (**A**) *fascin* mutant with global GAL4 (*fascin*^*sn28/sn28*^; *actin5c GAL4/+)*. (**B**) Global GFP-Fascin expression in *fascin* mutant (*fascin*^*sn28/sn28*^; *actin5c GAL4/UAS-GFP-Fascin)*. (**C**) Glocal GFP-Fascin-S52E expression in *fascin* mutant (*fascin*^*sn28/sn28*^; *actin5c GAL4/UAS-GFP-Fascin-S52E)*. (**D, E**) Graphs of quantification of pMRLC intensity at the nurse cell membranes (**D**) and border cell cluster (**E**) in the indicated genotypes. Each circle represents a follicle. Error bars=SD. ns indicates p>0.05, *p<0.05, **p<0.01 (One-way ANOVA with Tukey’s multiple comparison test). In **D**, peak pMRLC intensity was quantified at the nurse cell membranes and normalized to phalloidin staining in the same follicle, three measurements were taken per follicle and averaged. In **E**, pMRLC of intensity the border cell cluster was quantified and normalized to background staining in the same follicle. Restoring wild-type Fascin expression in both the somatic and germline cells of a *fascin* mutant follicle (B) significantly reduces activated Myosin enrichment on the nurse cell membranes (D) and border cell cluster (E) compared to the *fascin*-null control (B, D, E). Whereas expressing a phosphomimic form of Fascin in a *fascin* mutant (C) does not alter activated Myosin on the nurse cell membranes (D) or border cell cluster (E).

As we found that tight regulation of Myosin activity by Fascin is critical for on-time border cell migration (Figure 3), and expression of phosphomimetic Fascin (S52E) in *fascin* mutant follicles fails to restore normal levels of Myosin activity (Figure 4B-E), we expected it would also fail to fully rescue the delays in border cell migration. We previously found global expression (*actin 5c* GAL4) of wild-type Fascin in *fascin* mutant follicles rescues delays in border cell migration (Lamb et al., 2020). As expected, when we quantify the migration index for *fascin* mutant follicles with global expression of phosphomimietic Fascin (S52E), we find it only partially rescues delays in border cell migration (Figure 4-figure supplement 1C, migration index 0.90 compared to 0.80). Together these data indicate Fascin functions in other ways besides bundling actin and limiting Myosin activity to promote on-time border cell migration.

### Fascin acts in the border cells to control substrate stiffness

Previous evidence demonstrated that the stiffness of the nurse cells regulates border cell cluster stiffness as indicated by active Myosin levels and on-time border cell migration (Aranjuez et al., 2016). Since Fascin is required in both the nurse cells and the border cells to promote on-time border cell migration (Lamb et al., 2020), we wanted to determine which cells Fascin acts in to regulate Myosin activation. To test this, we used the UAS/GAL4 system to express a Fascin RNAi construct to knockdown Fascin in specific cell types – the germline (*matα* GAL4), somatic (*c355* GAL4) or the border cells (*c306* GAL4) – and analyzed how loss of Fascin in these different cells affects Myosin activation throughout the follicle. We have previously validated the use of UAS/GAL4 system to knockdown Fascin in these cell types (Lamb et al., 2020).

Based on the literature, we hypothesized knockdown of Fascin in the germline would increase Myosin activation in both the nurse cells and border cells, while knockdown of Fascin in the border cell would only increase Myosin activation in the border cells. We observe, as expected, knockdown of Fascin in the germline results in a significant increase in pMRLC (active MRLC) enrichment on the nurse cell membranes (Figure 5B, orange arrows and E, p<0.0001). However, knockdown down of Fascin in the germline unexpectedly fails to alter active MRLC enrichment on the border cell cluster (Figure 5B, blue arrow and F, p>0.05). We next knocked down Fascin in all the somatic cells or just the border cells and anticipated that this would lead to a significant increase in active MRLC on the border cell cluster but not the nurse cells. As expected, we observe a significant increase of active MRLC on the border cell cluster when Fascin is knocked down in the border cells (Figure 5C, D, F, blue arrows, p<0.0001). Surprisingly, knockdown of Fascin in the somatic or just the border cells also significantly increased active MRLC enrichment on the nurse cells (Figure 5C-E, orange arrows, p<0.0001). These data surprisingly suggest that knockdown of Fascin in the border cells increases border cell stiffness and this, in turn, induces the stiffening of their substrate, the nurse cells.

**Figure 5:**
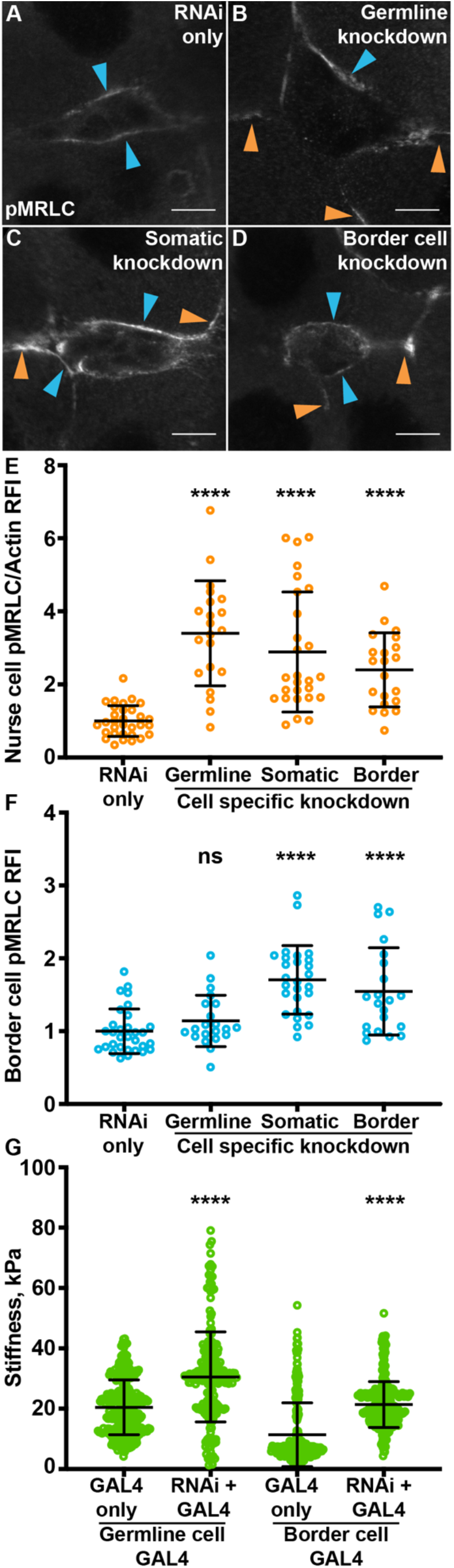
Germline Fascin knockdown increases myosin activation on the nurse cells while somatic Fascin knockdown increases myosin activation on both the border and nurse cells. (**A-D**) Maximum projections of 2-4 confocal slices of Stage 9 follicles of the indicated genotypes stained for phospho-MRLC (pMRLC, white). Blue arrows= pMRLC enrichment on border cell cluster. Orange arrows= pMRLC enrichment on surrounding nurse cells. Scale bars=10μm. (**A**) RNAi only (*fascin RNAi/+)*. (**B**) Germline knockdown of Fascin (*matα GAL4(3)/fascin RNAi*). **(C)**. Somatic cell knockdown of Fascin (*c355 GAL4/+; fascin RNAi*/+). (**D**) Border cell knockdown of Fascin (*c306 GAL4/+; fascin RNAi*/+). (**E, F**) Graphs of quantification of pMRLC intensity at the nurse cell membranes (**E**) and border cell cluster (**F**) in the indicated genotypes. Each circle represents a follicle. Error bars=SD. ns indicates p>0.05, ****p<0.0001 (One-way ANOVA with Tukey’s multiple comparison test). In **E**, peak pMRLC intensity was quantified at the nurse cell membranes and normalized to phalloidin staining in the same follicle, three measurements were taken per follicle and averaged. In **F**, pMRLC of intensity the border cell cluster was quantified and normalized to background staining in the same follicle. (**G**) Graph of nurse cell stiffness (kPa) of the indicated genotypes as measured by AFM. Each circle represents a single indentation. ****p<0.0001 (unpaired t-test). Error bars=SD. Fascin regulates myosin activation in the germline (B, E) and somatic cells (C, D, F). Loss of Fascin in the germline cells increases myosin activity and stiffness of the nurse cells (B, E, G). Loss of Fascin in the somatic or border cells increases myosin activity and stiffness of the nurse cells (B, E, G), and myosin activity in the border cell cluster (C, D, F).

Further, we used AFM to directly assess the changes in nurse cell stiffness of our cell specific Fascin knockdowns. The germline knockdown of Fascin results in nurse cells that are 1.5X stiffer than their GAL4 control (Figure 5G, p<0.0001), while the border cell knockdown results in nurse cell that are 1.8X stiffer than their GAL4 control (Figure 5G, p<0.0001). Together these data demonstrate the novel finding that Fascin acts the border cell cluster to regulate the stiffness of the surrounding nurse cell substrate (Figure 7).

### Fascin acts in the somatic cells to control Myosin activity throughout the follicle

Our RNAi experiments indicate that Fascin acts primarily in the border cells to control Myosin activation and nurse cell stiffness. If this is true, then restoring Fascin expression in only the somatic cells of a *fascin* mutant follicle, including the border cells, should restore normal Myosin activation in both the border cells and the nurse cells, and normal nurse cell stiffness. Indeed we find that expressing GFP-Fascin in the somatic cells of *fascin*-null follicles significantly reduces active MRLC enrichment on both the nurse cell and border cell membranes compared to the *fascin*-null control (Figure 6A-F, p<0.0001). Further, restoring Fascin expression in the somatic cells of *fascin*-null follicles significantly reduced the stiffness of the nurse cells compared to the *fascin*-null control (Figure 6G, 19.4 kPa compared to 38.9 kPa, p<0.0001). Together our data indicate that Fascin acts in the border cells to regulate the stiffness of both the border cell cluster and its substrate, the nurse cells.

**Figure 6:**
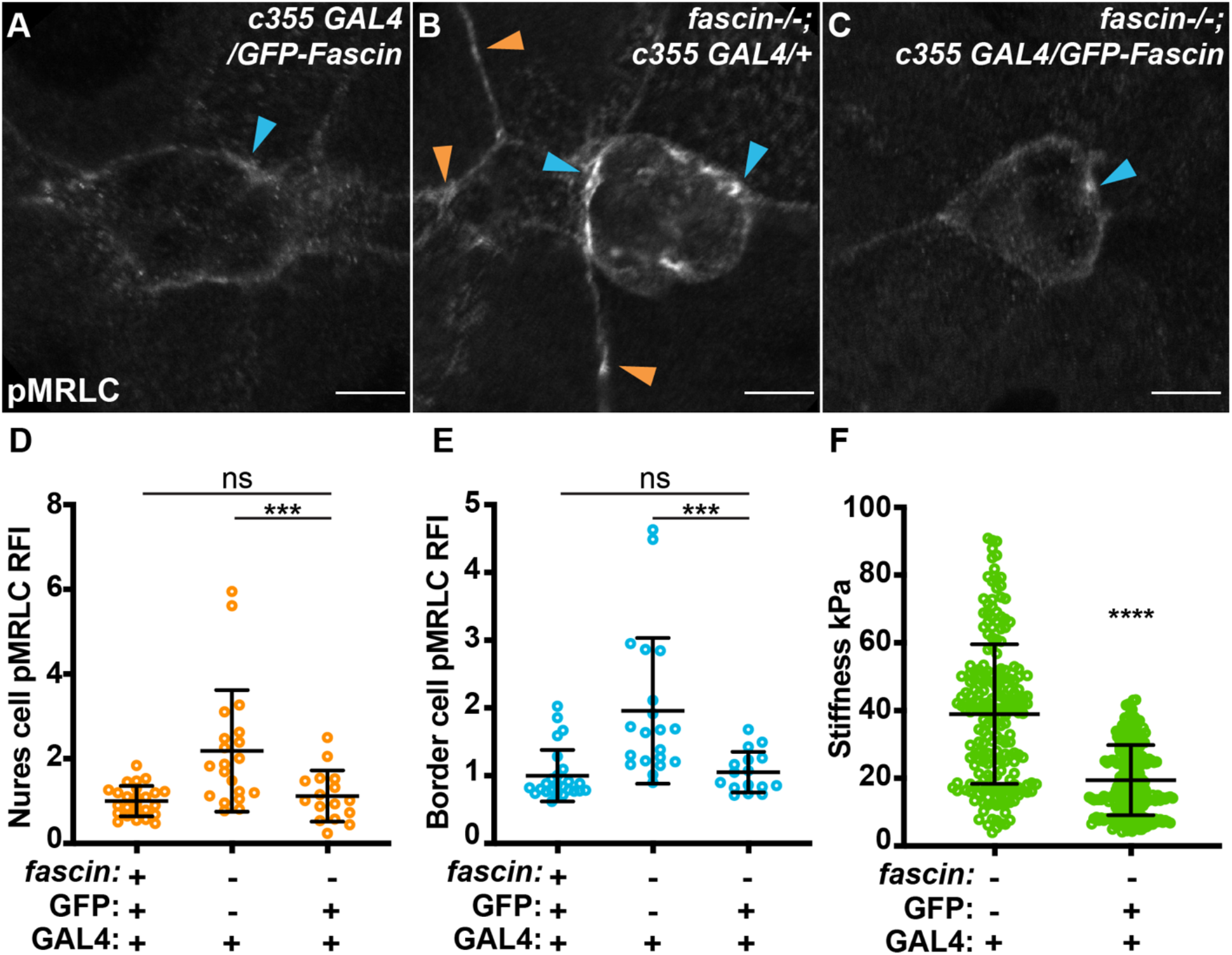
Somatic rescue of Fascin reduces nurse cell Myosin activity and stiffness. (**A-C**) Maximum projections of 2-4 confocal slices of Stage 9 follicles of the indicated genotypes stained for phospho-MRLC (pMRLC, white). Blue arrows= pMRLC enrichment on border cell cluster. Orange arrows= pMRLC enrichment on surrounding nurse cells. Scale bars=10μm. (**A**) Somatic GFP-Fascin expression (*c355 GAL4/+; UAS-GFP-Fascin/+)*. (**B**) *fascin* mutant with somatic GAL4 (*c355 GAL4, fascin*^*sn28/sn28*^). (**C**) Somatic GFP-Fascin expression in *fascin* mutant (*c355 GAL4, fascin*^*sn28/sn28*^; *UAS-GFP-Fascin/+)*. (**D, E**) Graphs of quantification of pMRLC intensity at the nurse cell membranes (**D**) and border cell cluster (**E**) in the indicated genotypes. Each circle represents a follicle. Error bars=SD. ns indicates p>0.05, ***p<0.0001 (One-way ANOVA with Tukey’s multiple comparison test). In **D**, peak pMRLC intensity was quantified at the nurse cell membranes and normalized to phalloidin staining in the same follicle, three measurements were taken per follicle and averaged. In **E**, pMRLC of intensity the border cell cluster was quantified and normalized to background staining in the same follicle. (**F**) Graph of nurse cell stiffness (kPa) of the indicated genotypes as measured by AFM. Each circle represents a single indentation. Error bars=SD. ****p<0.0001 (unpaired t-test). Restoring Fascin expression in the somatic cells of a *fascin* mutant follicle (C) significantly reduces activated Myosin enrichment on the nurse cell membranes (D) and border cell cluster (E) and reduces nurse cell stiffness by AFM (F) compared to the *fascin*-null control.

Given the surprising nature of our findings, we next wanted to determine if the border cell regulation of nurse cell stiffness is specific to Fascin or if it is a general principle. To test this idea, we expressed a constitutively active form of Rok (Rok-CAT) in the border cells (*c306* GAL4). Rok is one of the kinases that phosphorylates MRLC to activate Myosin. Thus, expressing constitutively active Rok will increase activation of Myosin, which, in turn, will increase cortical tension and therefore the stiffness of the border cells. We find that expression of constitutively active Rok in the border cells significantly increases active MRLC enrichment on both the nurse cell membranes (Figure 6-figure supplement 1B, orange arrows, and C, p<0.0001) and the border cell cluster (Figure 6-figure supplement 1B, blue arrows and D, p<0.001). These data suggest that the nurse cells, in general, respond to changes in stiffness of the border cells by altering their own cellular stiffness (Figure 7). This non-autonomous regulation of substrate stiffness by migratory cells is a novel and unexpected finding.

**Figure 7:**
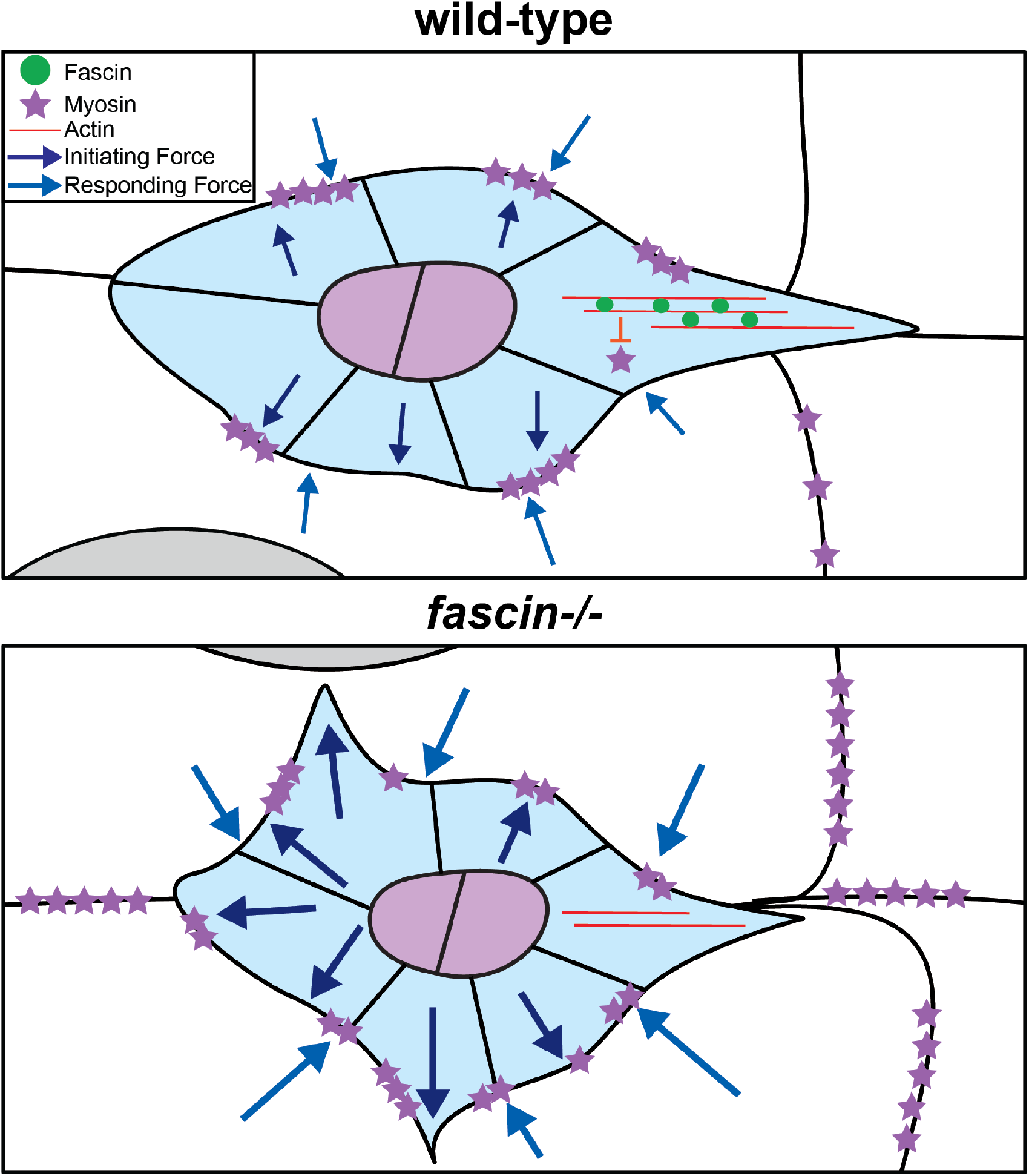
Proposed model for Fascin limiting Myosin activity to control substrate stiffness during border cell migration. In wild-type border cell clusters, Fascin (green circles) bundles actin (red lines) to limit Myosin activity (purple stars). Myosin activity in the border cell cluster generates forces (dark blue arrows) that pushes on the nurse cells which results in the nurse cells responding with force (lighter blue arrows). This balance of forces is required for on-time migration. In *fascin* mutant border cell clusters, Myosin activity on the border cell cluster is increased, driving increased Myosin activity on the nurse cells. This imbalance of forces between the border cell cluster and the nurse cell substrate impairs border cell migration.

## Discussion

Using *Drosophila* border cell migration as a model, we provide the first evidence that Fascin limits Myosin activity *in vivo* to control tissue stiffness (Figure 7). We find that loss of Fascin significantly increases activated Myosin, and this increase in Myosin activity contributes to the border cell migration delays observed in *fascin* mutant follicles during S9. Our data shows that Fascin bundling activity is required to limit Myosin activation, supporting the prior proposed model that Fascin tightly bundles actin and precludes Mysoin from binding to actin filaments (Elkhatib et al., 2014). The increased Myosin activation in *fascin* mutants results in substrate stiffening. Using cell-specific knockdown and rescue experiments, we made the suprising finding that Fascin activity in the border cells is necessary and sufficient to regulate Myosin activity and stiffness of the nurse cells. Thus, Fascin activity within the border cells plays a critical role in controlling the balance of forces between the border cells and their substrate, the nurse cells. We also show that this force balance is not specific to Fascin, as directly altering Myosin activity within the border cells phenocopies knockdown of Fascin in these cells. Together our data uncover the transformative finding that collectively migrating cells modulate the stiffness of their substrate (Figure 7).

Multiple lines of evidence support the model that Fascin is a critical regulator of cellular and tissue stiffness. Due to Myosin’s roles in contraction, readouts of Myosin activity can be used to indirectly assess relative cellular and tissue stiffnesses. We find that loss of Fascin causes increased Myosin activitation on both the border cell and nurse cell membranes (Figure 1). This finding suggests that loss of Fascin increases both the stiffness of the border cell cluster and its substrate, the nurse cells. Dynamic cycling of Myosin activity and inactivity is also essential for controlling stiffness and cell migration. Such dynamics can be indirectly assessed through live imaging of MRLC (Aranjuez et al., 2016; Majumder et al., 2012). We find that loss of Fascin results in slowed Myosin dynamics on the border cell cluster (Figure 1G-I, Video 1, 2). Additionally, loss of Fascin results in an increase in the number of active Myosin puncta (Figure 1E), consistent with our finding that there is increased Myosin activation at the border cell membranes. Notably, loss of Myosin activity in the border cells by RNAi knockdown of Rok or expression of a constitutively active form of Myosin phosphatase (Mbs) exhibits the opposite phenotype – decreased GFP-MRLC puncta (Aranjuez et al., 2016). Further, the active MRLC puncta on the border cells of *fascin* mutant follicles are also shorter in length, suggesting that loss of Fascin may also impact the distribution of active Myosin on the border cell cluster (Figure 1F). Together these two approaches of assessing Myosin activity support the model that Fascin plays critical roles in negatively regulating Myosin activitation in the border cells, and provides evidence that Fascin also functions to limit Myosin activity in the nurse cells. Thus, these data provide indirect cellular evidence that Fascin modulates the stiffness of both the border cells and nurse cells.

To directly assess tissue stiffness, we used AFM to determine the elastic moduli of S9 follicles. Based on the literature, we used two different probe indentation depths to measure the stiffness of the overlying basement membrane versus the underlying nurse cells (Figure 2; (Chlasta et al., 2017)). We find that basement membrane stiffness of *fascin* mutant follicles is not different from that of wild-type follicles (Figure 2D), indicating Fascin does not play a role in regulating the extracellular matrix surrounding the follicles. Two different groups have previously applied AFM to measure the stiffness of the basement membranes of various stages of *Drosophila* oogenesis (Chen et al., 2019; Chlasta et al., 2017; Crest et al., 2017). The stiffness values published by these two groups are very different. Specifically, Chlasta *et al*. published that S9 follicles have a range of stiffnesses from ∼100-400kPa across their anterior to posterior axis, with a mid-S9 follicle having a ∼300kPa basement membrane stiffness. Conversely, the other group found that S8 follicles have a stiffness ranging from ∼40-70kPa (Chen et al., 2019; Crest et al., 2017). One possible explanation for the different values that was proposed by Chlasta *et al*., was that the two groups performed a different number of measurements per follicle. Both groups used a range of 0-50nm, while we probed the entire depth of the basement membrane with an indentation range of 20-200nm. Overall, our basement membrane stiffness measurements of ∼25kPa are more similar to those seen by Chen *et al*. and Crest *et al*., and while there is some variability the ranges do overlap. Additionally, the minor differences in the stiffness values we observe may be due to the developmental stage assessed.

We used a deeper AFM probe indentation range to assess the stiffness of the nurse cells in our S9 follicles (Figure 2). Loss of Fascin results in a 2x increase in the nurse cell stiffness compared to wild-type follicles (Figure 2D). These data correlate well with the relative change in Myosin activation observed in the nurse cells of *fascin* mutant follicles (Figure 1). Further, we find that collagenase treatment, which degrades the basement membrane and allows nurse cell stiffness to be assessed at the surface of the follicle, reduces the stiffness measured at 20-200nm probe depth to be similar to that of the wild-type nurse cells (data not shown). Thus, we believe our AFM measurements accurately reflect the relative stiffness of the different layers within the S9 follicle, and allow us to assess changes in nurse cell stiffness due to genetic perturbations. Together our AFM and Myosin data indicate that Fascin plays a critical role in negatively regulating nurse cell stiffness.

We find that Fascin acts primarily within the border cells to control the stiffness of their substrate, the nurse cells. Cell-specific RNAi knockdown of Fascin reveals that while knockdown in the nurse cells does increase Myosin activation within the nurse cells, it fails to alter Myosin activation in the border cells. This finding was unexpected, as previous data in the field showed that increased Myosin activity in the nurse cells causes increased Myosin activation on the border cell cluster (Aranjuez et al., 2016). Specifically, this study overexpressed a Rho GEF in the nurse cells, which both increased Myosin activation and caused the nurse cells to change their shape and become more circular, ultimately impairing border cell migration (Aranjuez et al., 2016). As we do not observe any obvious changes in nurse cell shape when Fascin is lost or knocked down in the nurse cells, it may be that loss of Fascin does not cause a severe enough change in nurse cell Myosin activity and cell stiffness to cause the border cell cluster to respond. However, RNAi knockdown of Fascin in the nurse cells causes a ∼1.5x increase nurse cell stiffness in (Figure 5G). Conversely, RNAi knockdown of Fascin in either all somatic cells or the border cells increases Myosin activation in both the nurse cells and the border cells, and increases nurse cell stiffness ∼1.9x (Figure 5E-G). Further, knockdown of MRLC in the germline cells does not reduce activated Myosin on the border cell cluster (Figure 3-figure supplement 1D), highlighting the importance of Fascin in regulating Myosin in the border cell cluster. Similarly, restoring expression of Fascin in the somatic cells of *fascin*-null follicles restores both Myosin activity and nurse cell stiffness to wild-type levels (Figure 6). It is important to note that genetic background appears to affect nurse cell stiffness, as the germline and border cell GAL4 drivers have different baseline stiffnesses (Figure 5-6). Together, our data indicate that while Fascin does act within the nurse cells to regulate their stiffness, Fascin within the border cells is necessary and sufficient to control nurse cell Myosin activity levels and cell stiffness.

Fascin regulation of Myosin activity in the nurse cells is critical for border cell migration. We find that pharmacologically reducing Myosin activity or RNAi knockdown of MRLC in the germline of *fascin*-null follicles restores on-time border cell migration (Figure 3). These data support that Fascin-dependent inhibition of nurse cell Myosin activity and nurse cell stiffness is essential for on-time border cell migration. Further, our data indicates that this regulation of Myosin activation within the nurse cells is the result of Fascin’s activity within the border cells (Figure 5). We find that the phosphorylation state of Fascin regulates Myosin activation. Expression of the phosphomimetic form of Fascin (S52E), which is unable to bundle actin, in *fascin* mutants fails to both inhibit Myosin activation (Figure 4) or fully restore migration (Figure 4-figure supplement 1). Together, these data support the model that Fascin bundled actin precludes Myosin binding to actin and thereby, restricts its activation to promote the collective migration of the border cells.

Additionally, our finding that phosphomimetic Fascin partially rescues the migration delay in *fascin* mutants supports that non-bundling roles of Fascin also contribute to border cell migration. Indeed, Fascin has many functions besides actin bundling, such regulating microtubules (Villari et al., 2015) and acting within the nucleus (Groen et al., 2015). Additionally, S52 phosphorylated Fascin functions as an adaptor for the Linker of the Nucleoskeleton and Cytoskeleton (LINC) Complex (Groen et al., 2015; Jayo et al., 2016). This LINC Complex role of Fascin is required for nuclear shape changes necessary for mammalian single cell invasive migration (Jayo et al., 2016), raising the idea that Fascin may be similarly required for the invasion of the border cells between the nurse cells. Further experiments are needed to understand how the different functions of Fascin are coordinated to promote migration.

Together, our results suggest that increased stiffness in the border cell cluster affects the stiffness of its substrate, the nurse cells. This non-autonomous function of the border cells in altering the stiffness of the nurse cells is a novel observation. Previous data suggested that a balance of forces must be maintained between the border cells and the nurse cells, in which the nurse cells exert force on the border cells and the border cells respond to this force (Aranjuez et al., 2016). This balance of forces is considered necessary to promote the migration of the cluster through the tightly packed nurse cells (Aranjuez et al., 2016). Our data suggests that the border cells play a larger role in this balance of forces by exerting force on the nurse cells to control nurse cell stiffness. This interaction could potentially allow the border cell cluster to stiffen the nurse cells as the cluster migrates. Interestingly, in the context of cancer cell migration, a stiffer substrate often promotes cell migration (Oakes, 2018; Parekh and Weaver, 2016; Ren et al., 2021). Further, there is growing evidence that one means of directing migration is a gradient of substrate stiffness, such that cells move from softer to stiffer substrates; this is termed durotaxis (Shellard and Mayor, 2021; Sunyer and Trepat, 2020). Indeed, durotaxis has emerged as a property of collectively migrating cells. Specifically, it has been suggested that clusters of migrating cells are better able to sense differences in stiffness and respond more effectively (Martinez et al., 2016; Sunyer et al., 2016). Therefore, it is tempting to speculate that the border cells exert force on the nurse cells to stiffen them to aid in migration.

While it is clear that the balance of forces between the border cells and the nurse cells is critical for border cell migration, the mechanisms by which force imbalances impair migration remain poorly understood. We speculate that the increased Myosin activity in *fascin* mutants delays migration by impacting delamination and protrusion dynamics. We previously found that Fascin is required for on-time delamination of the border cell cluster from the follicular epithelium and for restricting the number and location of protrusions to the leading edge of the border cell cluster (Lamb et al., 2020). Similarly, both loss and constitutive activation of Myosin within the border cells delays delamination, and causes excessive and misdirected protrusions (Aranjuez et al., 2016; Majumder et al., 2012; Mishra et al., 2019). These data suggest that it is not only the level of Myosin activity, but is ability to cycle between active and inactive states that contributes to these two aspects of border cell migration. Based on our data, we suspect altering Myosin activity in the border cells ultimately changes the stiffness of the nurse cells. Too little activation of Myosin would result in a soft substrate and too much would result in a stiff substrate. Such changes in stiffness could alter the polarization of the cluster, resulting in mislocalized and increased protrusions which not only delay migration but impair delamination. Supporting this idea, Myosin regulates active Rac polarization within the border cells (Mishra et al., 2019). Rac activation is highest in the leading cell of the border cell cluster and is require to generate forward directed protrusions (Bianco et al., 2007; Fulga and Rorth, 2002; Mishra et al., 2019). Increased Myosin activation in the border cell cluster disrupts this polarization, resulting in mislocalized protrusions (Mishra et al., 2019). This loss of polarization could function cell-autonomously, but, based on our data, it may also increase nurse cell stiffness. Such increased stiffness could impair delamination and cause mislocalized protrusions by physically altering the topography of the nurse cells, which has recently been shown to be critical for border cell migration and forward directed protrusions (Dai et al., 2020). Additionally, increased substrate stiffness could disrupt durotactic signaling or alter the diffusion of the ligands directing migration. Thus, Fascin limiting Myosin activation likely contributes to the delayed delamination and aberrant mislocalized protrusions observed during border cell migration in *fascin*-null follicles.

Our discovery that Fascin limits Myosin activity *in vivo* is a novel finding that is unlikely to be restricted to *Drosophila*. Indeed, both Fascin and Myosin play critical roles during cancer metastasis (Aguilar-Cuenca et al., 2014; Hashimoto et al., 2011; Ma and Machesky, 2015). Increased Myosin activation and consequently, increased stiffness are a common phenotype observed in cancer cells and their substrate (Aguilar-Cuenca et al., 2014; Ren et al., 2021; Tse et al., 2012; van Helvert and Friedl, 2016). Increased substrate stiffness promotes migration in a wide range of cancers, suggesting increased Myosin activity can lead to increased cancer metastasis (Aguilar-Cuenca et al., 2014; Emon et al., 2018; Mierke, 2020; Ren et al., 2021). Additionally, Fascin is highly expressed in many types of cancers, notably carcinomas (Hashimoto et al., 2011; Ma and Machesky, 2015). High Fascin expression in these cancers is correlated with increased migration (Grothey et al., 2000; Hashimoto et al., 2007), invasion (Adams et al., 1999; Minn et al., 2005), and metastasis (Alburquerque-Gonzalez et al., 2020; Li et al., 2014), highlighting Fascin as a critical promoter of cancer cell migration. However, according to our model, increased Fascin would limit Myosin activity. Given the large focus on Fascin as an actin bundling protein and our finding that Fascin-dependent bundling is required to limit Myosin activity and substrate stiffness suggests that phosphorylated Fascin may promote cancer metastasis by allowing high Myosin activation and potentially other bundling-independent functions. Supporting this idea, expression of a S39 phosphomimetic form of Fascin, which can not bundle actin, promotes human colon carcinoma migration (Hashimoto et al., 2007), suggesting phosphorylated Fascin could promote cancer metastasis by allowing increased Myosin activation and cell stiffness.

The mechanical communication between migrating cells and their substrate is a growing area of research. Until now, the premise in the field has been that substrate stiffness regulates the mechanical properties of the migrating cells and thereby, alters their ability to migrate. For example, in a model of breast cancer cell migration, high substrate stiffness promotes migration (Ren et al., 2021). Additionally during zebrafish development, the underlying mesoderm must stiffen to induce the epithelial to mesenchymal transition (EMT) and migration of the neural crest cells (Barriga et al., 2018). Together these studies highlight the current paradigm that substrate stiffness is the driving force that regulates the migrating cells to control their migration. Here we purpose a paradigm shifting interaction, in which the stiffness of the migrating cells regulates substrate stiffness to promote migration. Our transformative findings suggest that during collective cell migrations, such as those during development and cancer metastasis, the migrating cells apply force to induces the stiffening of their substrate, this results in a reciprocal mechanical communication between the migrating cells and their substrate. This concept is consistent with the idea of migrating cells altering their ECM environment by changing its composition or structure to promote their migration. Overall, our findings shift the paradigm in the field from the substrate controlling migrating cell stiffness and thereby, migration, to the migrating cells also being able to alter their environment and substrate stiffness to promote their own migration.

## Figure legends

**Video 1: Myosin dynamics in control follicle**. Video of S9 control follicle (*fascin*^*sn28*^/+; *GFP-MRLC*/+). Time listed in seconds. Images were acquired every 30 seconds with a 20x objective. Anterior is to the right. Scale bar = 20μm. The control cluster displays Myosin dynamics in which Myosin puncta appear and disappear rapidly on the border cell cluster.

**Video 2: Myosin dynamics in *fascin*-null follicle**. Video of S9 control follicle (*fascin*^*sn28/sn28*^; *GFP-MRLC*/+). Time listed in seconds. Images were acquired every 30 seconds with a 20x objective. Anterior is to the right. Scale bar = 20μm. The control cluster displays Myosin dynamics in which Myosin puncta appear and disappear rapidly on the border cell cluster.

## Materials and Methods

### Fly stocks

Fly stocks were maintained on cornmeal/agar/yeast food at 21°C, except where noted. Before immunofluorescence and live imaging, flies were fed wet yeast paste daily for 2-4 days. Unless otherwise noted, *yw* was used as the wild-type control. The following stocks were obtained from the Bloomington Stock Center (Bloomington, IN): *mat α* GAL4 (third chromosome), *c355* GAL4, *c306* GAL4, *actin5C* GAL4, *UASp-RNAi-Fascin* (TRiP.HMS02450), *UASp-Sqh-RNAi* (TRiP.HMS00437), *UASp-Rok-CAT*. The *fTRG sqh* stock was obtained from the Vienna *Drosophila* Resource Center. The *fascin*^*sn28*^ line was a generous gift form Jennifer Zanet (Université de Toulouse, Toulouse, France (Zanet et al., 2012)), the *oskar* GAL4 line (second chromosome) was a generous gift from Anne Ephrussi (European Molecular Biology Laboratory, Heidelber, Germany (Telley et al., 2012)), the *UASp-GFP-Fascin* and *UASp-GFP-Fascin-S52E* lines were a generous gift from Francois Payre (Université de Toulouse, Toulouse, France (Zanet et al., 2009). For germline expression during S9, either *matα* GAL4 or *oskar* GAL4 were utilized interchangeably. Expression of *UASp-RNAi-Fascin* was achieved by crossing to *matα* GAL4, *c355* GAL4, and *c306* GAL4, maintaining crosses at 25°C and progeny at 29°C for 3 days. Expression of *UASp-Sqh-RNAi* was achieved by crossing to *oskar* GAL4, maintaining crosses at 25°C and progeny at 29°C for 3 days. The *sn28, c355* GAL4 flies were generated previously (Lamb et al., 2020). Expression of *UASp-GFP-Fascin* or *UASp-GFP-Fascin-S52E* was achieved by crossing to *actin5C* GAL4, crosses were maintained at 25°C and progeny at 29°C for 2 days.

### Immunofluorescence

Whole-mount *Drosophila* ovary samples (approximately 5 flies per experiment) were dissected into Grace’s insect media (Lonza, Walkersville, MD) and fixed for 10 minutes at room temperature in 4% paraformaldehyde in Grace’s insect media. Briefly, samples were blocked using Triton antibody wash (1X phosphate-buffered saline, 0.1% Triton X-100, and 0.1% bovine serum albumin) six times for 10 minutes each. Primary antibodies were diluted with Triton antibody wash and incubated overnight at 4°C. The following primary antibodies were obtained from the Developmental Studies Hybridoma Bank (DSHB) developed under the auspices of the National Institute of Child Health and Human Development and maintained by the Department of Biology, University of Iowa (Iowa City, IA): mouse anti-Hts 1:50 (1B1, Lipshitz, HD (Zaccai and Lipshitz, 1996)), mouse anti-FasIII 1:50 (7G10, Goodman, C (Patel et al., 1987)); mouse anti-Fascin 1:20 (sn7c, Cooley, L (Cant et al., 1994)). Additionally, the following primary antibody was used: rabbit anti-GFP 1:2000 (pre-absorbed on *yw* ovaries at 1:20 and used at 1:100; Torrey Pines Biolabs, Inc., Secaucus, NJ). After 6 washes in Triton antibody wash (10 minutes each), secondary antibodies were incubated overnight at 4°C or for ∼4 hours at room temperature. The following secondary antibodies were used at 1:500: AlexaFluor (AF)488::goat anti-mouse, AF568::goat anti-mouse, AF488::goat anti-rabbit, AF568::goat anti-rabbit (Thermo Fischer Scientific). AF647-, or AF568-conjugated phalloidin (Thermo Fischer Scientific) was included with primary and secondary antibodies at a concentration of 1:250. After 6 washes in Triton antibody wash (10 minutes each), 4’,6-diamidino-2-phenylindole (DAPI, 5 mg/ml) staining was performed at a concentration of 1:5000 in 1X PBS for 10 minutes at room temperature. Ovaries were mounted in 1 mg/ml phenylenediamine in 50% glycerol, pH 9 (Platt and Michael, 1983). All experiments were performed a minimum of three independent times.

Active-MRLC staining was performed using a protocol provided by Jocelyn McDonald (Majumder et al. 2012; Aranjuez et al. 2016). Briefly, ovaries were fixed for 20 min at room temperature in 8% paraformaldehyde in 1X phosphate-buffered saline (PBS) and 0.5% Triton X-100. Samples were blocked by incubating in Triton antibody wash (1XPBS, 0.5% Triton X-100, and 5% bovine serum albumin) for 30 min. Primary antibodies were incubated for 48 hr at 4°. The rabbit anti-pMRLC (S19; Cell Signaling, Davers, MA) was diluted 1:100 in Triton antibody wash, and anti-Fascin (sn7c, 1:20) was sometimes added to the primary antibody solution to differentiate between wild-type and *fascin*-null follicles in the same sample or to confirm Fascin RNAi knockdown. After six washes in Triton antibody wash (10 min each), the secondary antibodies were diluted 1:500 in Triton antibody wash and incubated overnight at 4°. Alexa Fluor 647–phalloidin (Invitrogen, Life Technologies, Grand Island, NY) was included with both primary and secondary antibodies at a concentration of 1:250. Samples were washed six times in Triton antibody wash (10 min each) and the stained with DAPI and mounted as described above.

### Image acquisition and processing

Microscope images of fixed *Drosophila* follicles were obtained using LAS AS SPE Core software on a Leica TCS SPE mounted on a Leica DM2500 using an ACS APO 20x/0.60 IMM CORR -/D objective (Leica Microsystems, Buffalo Grove, IL) or using Zen software on a Zeiss 700 LSM mounted on an Axio Observer.Z1 using a Plan-Apochromat 20x/0.8 working distance (WD) = 0.55 M27 or a EC-Plan-Neo-Fluar 40x/1.3 oil objective (Carl Zeiss Microscopy, Thornwood, NY). Maximum projections (two to four confocal slices), merged images, rotations, and cropping were performed using ImageJ software (Abramoff et al., 2004). S9 follicles were identified during fixed imaging by the size of the follicle (∼150-250μm), the position and morphology of the outer follicle cells, and presence of a border cell cluster. The beginning of S10 was defined as when the anterior most outer follicle cells reached the nurse cell-oocyte boundary and flattened.

### Quantification of fixed imaging for border cell migration

Quantification of the migration index of border cell migration was performed as described in (Fox et al., 2020; Lamb et al., 2020). Briefly, quantification of S9 follicles was performed on confocal image stacks of follicles stained with anti-Hts and anti-FasIII or phalloidin. Measurements of migration distances were obtained from maximum projections of 2-4 confocal slices of deidentified 20x confocal images using ImageJ software (Abramoff et al., 2004). Briefly, a line segment was drawn from the anterior end of the follicle to the front or posterior of the border cell cluster and the distance in microns measured, this was defined as the distance of border cell migration. Additionally, a line segment was drawn from the anterior end of the follicle to the anterior end of the main-body follicle cells and the distance measured, this was defined as the distance of the outer follicle cells. Lastly, the entire follicle length was measured along the anterior-posterior axis. The migration index was calculated in Excel (Microsoft, Redmond, WA) by dividing the border cell distance by the follicle cell distance. Cluster length was determined by measuring the distance from the front to the rear of the border cell cluster (detached cells were not included). Data was compiled, graphs generated, and statistical analysis performed using Prism (GraphPad Software).

### pMRLC quantifications

Intensity analysis were performed on maximum projections of 3 confocal slices of 40x confocal images using ImageJ software. For nurse cell intensity, 3 line segments per follicle were drawn across nurse cell-nurse cell membranes on maximum projections of 2-3 confocal slices of follicles stained for pMRLC and phalloidin. The fluorescent intensity peak for pMRLC was determined for each line and normalized to phalloidin intensity at the same point. These three values were then averaged for a single image. Averages were then normalized to the wild-type average for each experiment due to experimental variability. For border cell intensity, the border cell cluster was traced using the phalloidin stain and the mean fluorescence intensity for pMRLC was measured for this shape and this was then normalized to the mean fluorescence intensity of pMRLC of the same shape in the nurse cell cytoplasm. For the puncta number and length, puncta on the border cell cluster were manually counted and length measured from a max projection image using ImageJ software. Data was compiled, graphs generated, and statistical analysis performed using Prism (GraphPad Software).

### Live imaging

Whole ovaries were dissected from flies fed wet yeast past for 2-3 days and maintained at 25°C until the last 16-24 hours when they were moved to 29°C. Genotypes used for live imaging were *sn28/FM7; sqh-GFP*, or *sn28/sn28; sqh-GFP*. Ovaries were dissected in Stage 9 (S9) medium (Prasad *et al*. 2007): Schneider’s medium (Life Technologies), 0.6x penicillin/streptomycin (Life Technologies), 0.2 mg/ml insulin (Sigma-Aldrich, St. Louis, MO), and 15% fetal bovine serum (Atlanta Biologicals, Flowery Branch, GA). S9 follicles were hand dissected and embedded in 1.25% low-melt agarose (IBI Scientific, Peosta, IA) made with S9 media on a coverslip-bottom dish (MatTek, Ashland, MA). Just prior to live imaging, fresh S9 media was added to coverslip-bottom dish. Live imaging was performed with Zen software on a Zeiss 700 LSM mounted on an Axio Observer.Z1 using a Plan-Apochromat 20x/0.8 working distance (WD) = 0.55 M27 (Carl Zeiss Microscopy, Thornwood, NY). Images were acquired every 30 seconds for at least 1 hour for *Sqh-GFP* flies. Maximum projections (2-5 confocal slices), merge images, rotations, and cropping were performed using ImageJ software (Abramoff et al., 2004) To aid in visualization live imaging videos were brightened by 50% in Photoshop (Adobe, San Jose, CA).

### Quantification of live imaging

Quantification of live imaging videos were performed in ImageJ (Abramoff et al., 2004) using maximum projection of 2-5 confocal slices from time-lapse videos of border cell migration. For Sqh-GFP live imaging, puncta lifetime was defined by the amount of time elapsed from when a punctum first appeared to when it disappeared completely. Data were compiled, graphs generated, and statistical analysis performed using Prism (GraphPad Software).

### Atomic force microscopy (AFM) nanoindentation on *Drosophila* follicles

Whole ovaries were dissected from flies fed wet yeast past for 2-3 days. Ovaries were dissected in Stage 9 (S9) medium (Prasad *et al*. 2007), as described above. S9 follicles were hand isolated and mounted on poly-D-lysine coated 35 mm round glass coverslips. Force spectroscopy data were collected using a molecular force probe 3D (Asylum research) AFM in a liquid cell. AFM force spectroscopy was performed in a buffered solution within 1 – 2 hours after submersion. A new silicon nitride AFM probe (Bruker, DNP-10) was used for every experiment with a nominal spring constant of 0.12 N/m and a half cone angle of 20 degrees. Actual spring constant was calibrated using the built-in thermal noise method prior to measurement collection in each experiment. S9 follicles were located using the top view video camera and AFM force versus indentation data were collected on the middle of the follicle. The force data were recorded with a 0.6 – 1.2 μm/s tip approach velocity and a maximum force ranging from 1 – 5 nN. Typically, 2 – 3 different follicles were probed for each sample. In each region, 5 – 10 different positions with 2 – 10 μm separations were probed. For each position 3 – 8 multiple repeated force curves were recorded. Two stiffness values of follicles were determined by fitting tha approach data of two separate tip depth force-indentation curves to the rearranged form of the Hertzian elastic contact model (Heinrich, 1882). The two regions were selected to measure the stiffness of the basement membrane (20-200nm) and underlying nurse cells (200-800nm) and are similar to previous studies(Chen et al., 2019; Chlasta et al., 2017; Crest et al., 2017). Poisson’s ratios of 0.5 and 0.25 were assumed for the follicles and AFM probe, respectively. The data analysis was carried out as in our previously reported work (Bell et al., 2020; Kruger et al., 2019, 2020; McGowan et al., 2020).

### Pharmacological inhibition of Myosin in *Drosophila* follicles

Whole ovaries were dissected from flies fed wet yeast past for 2-3 days and maintained at room temperature. Ovaries of wild-type (*yw*) or *fascin* mutant (*fascin*^*sn28/sn28*^) flies were dissected in Stage 9 (S9) medium (Prasad *et al*. 2007), as described above. Ovarioles were teased apart and then were incubated at room temperature for 2 hours in either control media (S9 media + vehicle (DMSO)), 200µM of Y-27632, or 200µM of blebbistatin. After 2 hours, ovaries were rinsed 3 times with S9 media and then fixed and stained following the pMRLC staining protocol described above.

## Acknowledgments

We thank the Westside Fly Group and Dunnwald lab for helpful discussions, and the Tootle lab for helpful discussions and careful review of the manuscript. Stocks obtained from the Bloomington Drosophila Stock Center (NIH P40OD018537) were used in this study. Information Technology Services – Research Services provided data storage support. This project is supported by National Institutes of Health R01GM116885. M.C.L. was partially supported by the University of Iowa Summer Graduate Fellowship.

---

**Figure 3- figure supplement 1:**
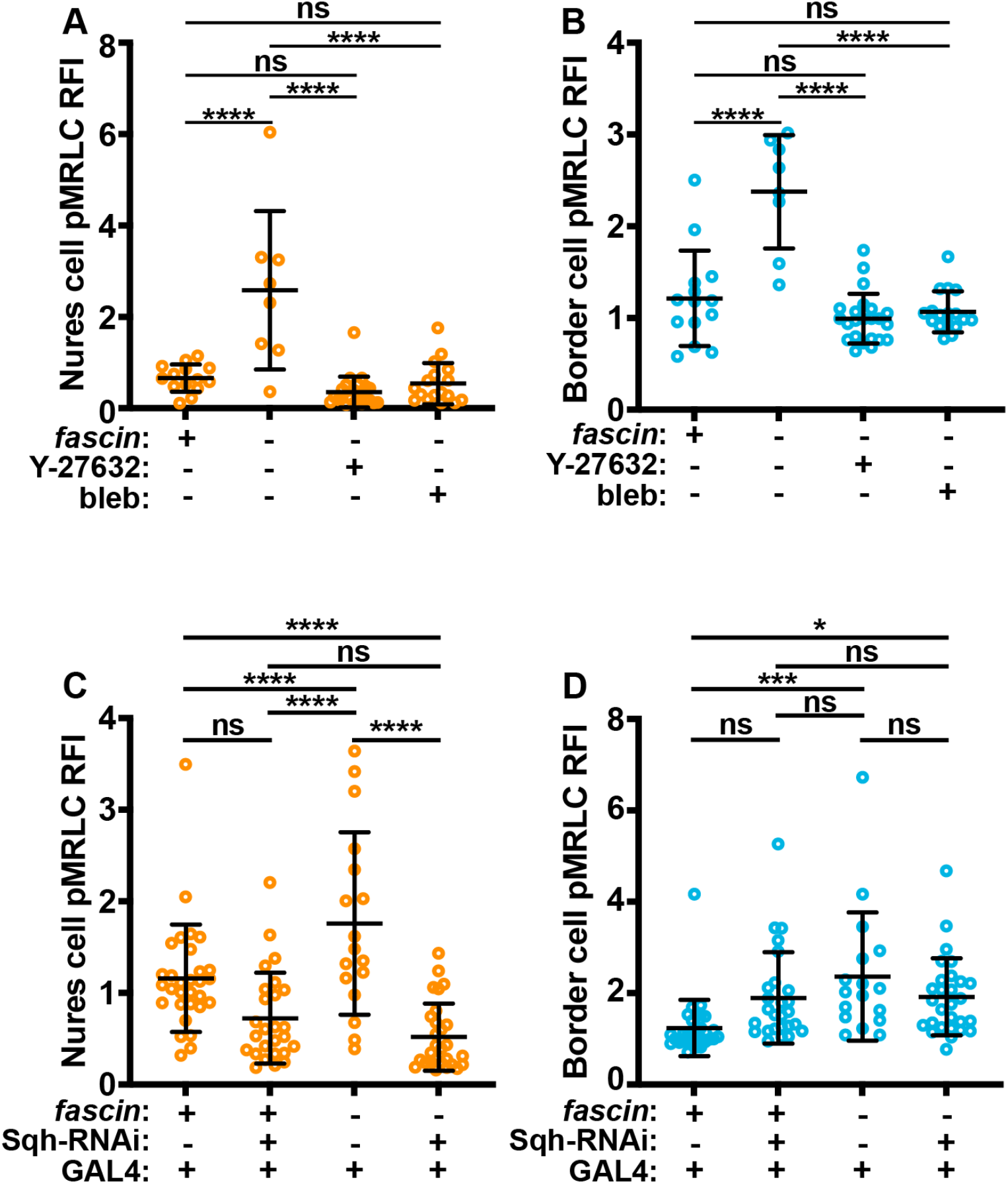
Pharmacological inhibition of Myosin and germline MRLC knockdown reduce active Myosin in the follicle. (**A-D**) Graphs of quantification of pMRLC intensity at the nurse cell membranes (**A, C**) and border cell cluster (**B, D**) in the indicated genotypes. Each circle represents a follicle. Error bars=SD. ns indicates p>0.05, *p<0.05, ***p<0.001, ****p<0.0001 (One-way ANOVA with Tukey’s multiple comparison test). In **A, C**, peak pMRLC intensity was quantified at the nurse cell membranes and normalized to phalloidin staining in the same follicle, three measurements were taken per follicle and averaged. In **B, D**, pMRLC of intensity the border cell cluster was quantified and normalized to background staining in the same follicle. Pharmacological inhibition of Myosin reduces active Myosin in the border cells and nurse cells (A, B), while germline knockdown of MRLC reduced activated Myosin on only the nurse cell membranes (C, D).

**Figure 4- figure supplement 1:**
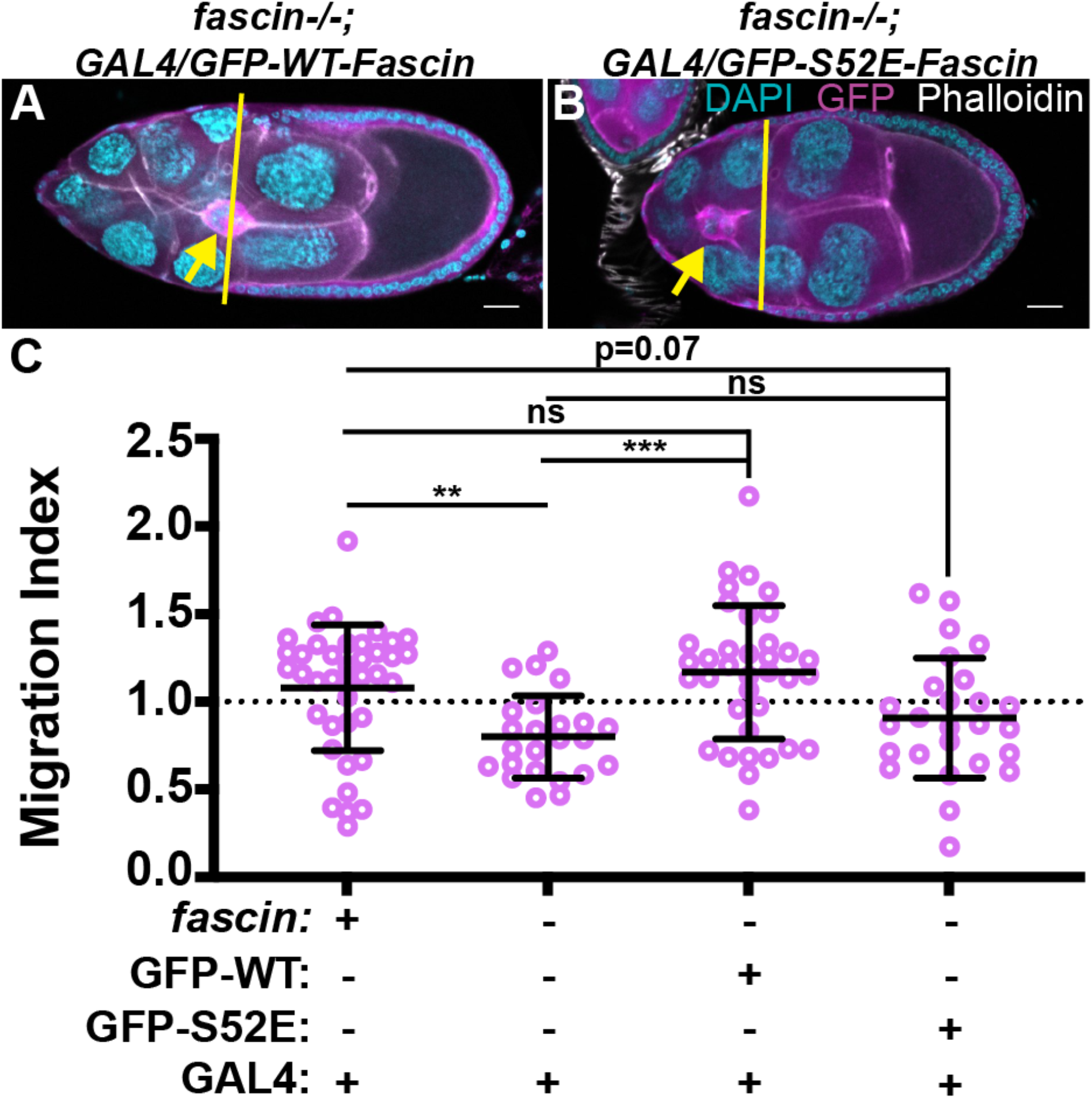
Phosphorylation of Fascin regulates border cell migration. (**A, B**) Maximum projections of 2-4 confocal slices of Stage 9 follicles of the indicated genotypes. Merged images: GFP-Fascin (magenta), phalloidin (white), and DAPI (cyan). Yellow lines = outer follicle cell distance. Yellow arrows = border cell cluster. Scale bars = 20 μm. (**A**) Global GFP-Fascin expression in *fascin* mutant (*fascin*^*sn28/sn28*^; *actin5c GAL4/UAS-GFP-Fascin)*. **(B)** Glocal GFP-Fascin-S52E expression in *fascin* mutant (*fascin*^*sn28/sn28*^; *actin5c GAL4/UAS-GFP-Fascin-S52E)*. (**C**) Migration index quantification of the indicated genotypes. Dotted line at 1 = on-time migration. Circle = S9 follicle. Lines = averages and error bars = SD. ns indicates p>0.05, **p < 0.01, ***p<0.001 (One-way ANOVA with Tukey’s multiple comparison test). Phosphomimetic Fascin expression in *fascin* mutants partially rescues delays in S9 border cell migration delay (A-C).

**Figure 6- figure supplement 1:**
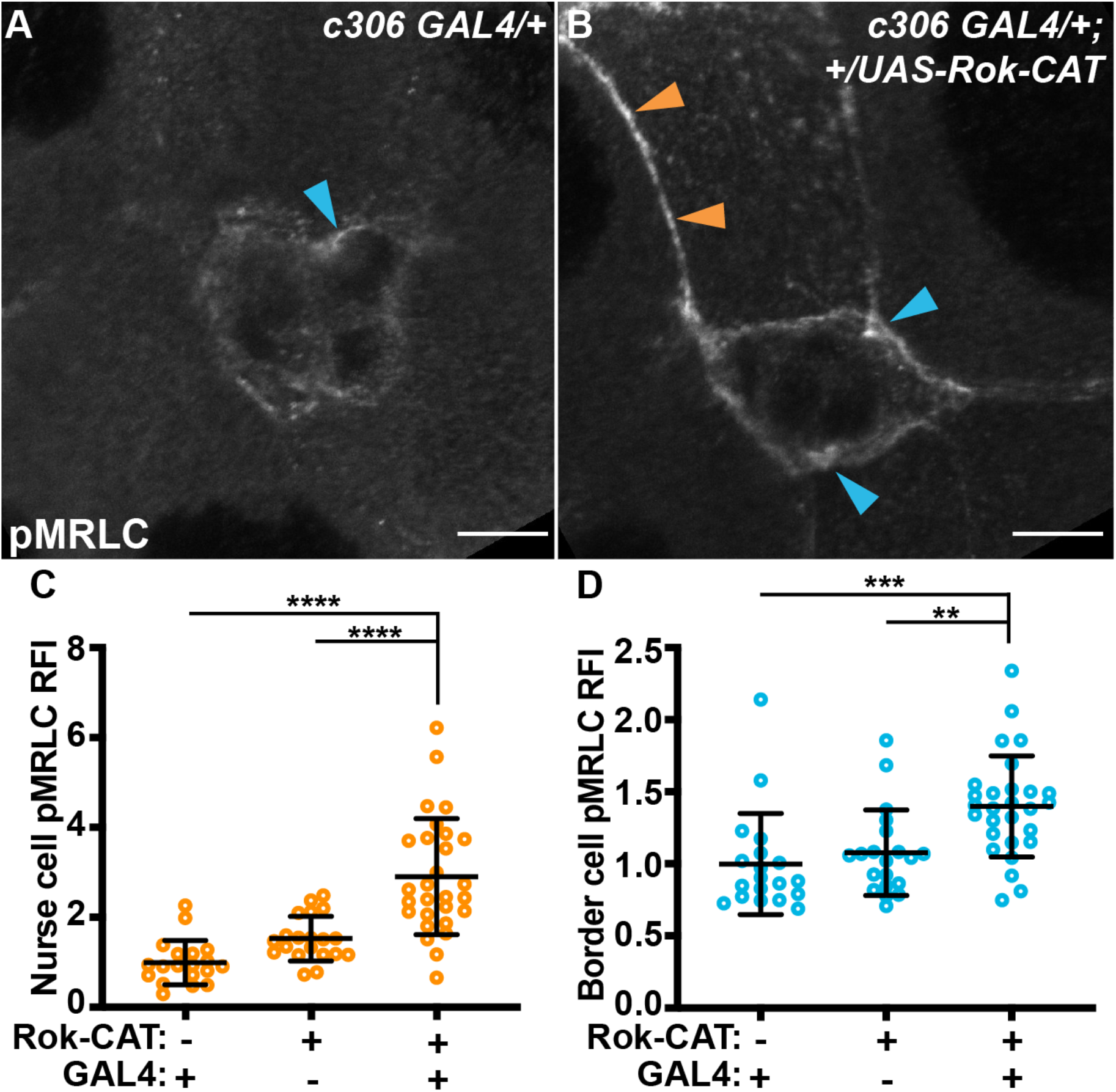
Increasing border cell stiffness through activated Rok increases activated Myosin on the nurse cells. (**A, B**) Maximum projections of 2-4 confocal slices of Stage 9 follicles of the indicated genotypes stained for phospho-MRLC (pMRLC, white). Blue arrows= pMRLC enrichment on border cell cluster. Orange arrows= pMRLC enrichment on surrounding nurse cells. Scale bars=10μm. (**A**) Border cell GAL4 only control (*c306 GAL4/+*). (**B**) Border cell expression of constitutively active Rok (*c306 GAL4/+; UAS-Rok-CAT/+*). (**C, D**) Graphs of quantification of pMRLC intensity at the nurse cell membranes (**C**) and border cell cluster (**D**) in the indicated genotypes. Each circle represents a follicle. Error bars=SD. **p<0.01, ***p<0.001 ****p<0.0001 (One-way ANOVA with Tukey’s multiple comparison test). In **C**, peak pMRLC intensity was quantified at the nurse cell membranes and normalized to phalloidin staining in the same follicle, three measurements were taken per follicle and averaged. In **D**, pMRLC of intensity the border cell cluster was quantified and normalized to background staining in the same follicle. Expression of constitutively active Rok in the border cells (B) leads to significantly increased activated Myosin enrichment on both the nurse cell membranes (C) and border cell cluster (D) compared to the GAL4 only control (A, C, D).

## References

Abramoff, M.D., Magalhaes, P., Ram, S., 2004. Image processing with ImageJ. Biophotonics Int 11, 36–42.

Adams, J.C., Clelland, J.D., Collett, G.D., Matsumura, F., Yamashiro, S., Zhang, L., 1999. Cell-matrix adhesions differentially regulate fascin phosphorylation. Mol Biol Cell 10, 4177–4190.

Aguilar-Cuenca, R., Juanes-Garcia, A., Vicente-Manzanares, M., 2014. Myosin II in mechanotransduction: master and commander of cell migration, morphogenesis, and cancer. Cell Mol Life Sci 71, 479–492.

Alburquerque-Gonzalez, B., Bernabe-Garcia, M., Montoro-Garcia, S., Bernabe-Garcia, A., Rodrigues, P.C., Ruiz Sanz, J., Lopez-Calderon, F.F., Luque, I., Nicolas, F.J., Cayuela, M.L., Salo, T., Perez-Sanchez, H., Conesa-Zamora, P., 2020. New role of the antidepressant imipramine as a Fascin1 inhibitor in colorectal cancer cells. Exp Mol Med 52, 281–292.

Aranjuez, G., Burtscher, A., Sawant, K., Majumder, P., McDonald, J.A., 2016. Dynamic myosin activation promotes collective morphology and migration by locally balancing oppositional forces from surrounding tissue. Mol Biol Cell 27, 1898–1910.

Barriga, E.H., Franze, K., Charras, G., Mayor, R., 2018. Tissue stiffening coordinates morphogenesis by triggering collective cell migration in vivo. Nature 554, 523–527.

Bell, K.J., Lansakara, T.I., Crawford, R., Monroe, T.B., Tivanski, A.V., Salem, A.K., Stevens, L.L., 2020. Mechanical cues protect against silica nanoparticle exposure in SH-SY5Y neuroblastoma. Toxicol in Vitro, 105031.

Bianco, A., Poukkula, M., Cliffe, A., Mathieu, J., Luque, C.M., Fulga, T.A., Rorth, P., 2007. Two distinct modes of guidance signalling during collective migration of border cells. Nature 448, 362–365.

Butcher, D.T., Alliston, T., Weaver, V.M., 2009. A tense situation: forcing tumour progression. Nat Rev Cancer 9, 108–122.

Cant, K., Knowles, B.A., Mooseker, M.S., Cooley, L., 1994. Drosophila singed, a fascin homolog, is required for actin bundle formation during oogenesis and bristle extension. J Cell Biol 125, 369–380.

Chanet, S., Miller, C.J., Vaishnav, E.D., Ermentrout, B., Davidson, L.A., Martin, A.C., 2017. Actomyosin meshwork mechanosensing enables tissue shape to orient cell force. Nat Commun 8, 15014.

Chang, S.S., Rape, A.D., Wong, S.A., Guo, W.H., Wang, Y.L., 2019. Migration regulates cellular mechanical states. Mol Biol Cell 30, 3104–3111.

Chen, D.Y., Crest, J., Streichan, S.J., Bilder, D., 2019. Extracellular matrix stiffness cues junctional remodeling for 3D tissue elongation. Nat Commun 10, 3339.

Chlasta, J., Milani, P., Runel, G., Duteyrat, J.L., Arias, L., Lamire, L.A., Boudaoud, A., Grammont, M., 2017. Variations in basement membrane mechanics are linked to epithelial morphogenesis. Development 144, 4350–4362.

Coravos, J.S., Mason, F.M., Martin, A.C., 2017. Actomyosin Pulsing in Tissue Integrity Maintenance during Morphogenesis. Trends Cell Biol 27, 276–283.

Crest, J., Diz-Munoz, A., Chen, D.Y., Fletcher, D.A., Bilder, D., 2017. Organ sculpting by patterned extracellular matrix stiffness. Elife.

Dai, W., Guo, X., Cao, Y., Mondo, J.A., Campanale, J.P., Montell, B.J., Burrous, H., Streichan, S., Gov, N., Rappel, W.J., Montell, D.J., 2020. Tissue topography steers migrating Drosophila border cells. Science 370, 987–990.

De Pascalis, C., Etienne-Manneville, S., 2017. Single and collective cell migration: the mechanics of adhesions. Mol Biol Cell 28, 1833–1846.

Di Martino, J., Henriet, E., Ezzoukhry, Z., Goetz, J.G., Moreau, V., Saltel, F., 2016. The microenvironment controls invadosome plasticity. J Cell Sci 129, 1759–1768.

Eble, J.A., Niland, S., 2019. The extracellular matrix in tumor progression and metastasis. Clin Exp Metastasis 36, 171–198.

Edwards, K.A., Kiehart, D.P., 1996. Drosophila nonmuscle myosin II has multiple essential roles in imaginal disc and egg chamber morphogenesis. Development 122, 1499–1511.

Elkhatib, N., Neu, M.B., Zensen, C., Schmoller, K.M., Louvard, D., Bausch, A.R., Betz, T., Vignjevic, D.M., 2014. Fascin plays a role in stress fiber organization and focal adhesion disassembly. Curr Biol 24, 1492–1499.

Emon, B., Bauer, J., Jain, Y., Jung, B., Saif, T., 2018. Biophysics of Tumor Microenvironment and Cancer Metastasis - A Mini Review. Comput Struct Biotechnol J 16, 279–287.

Fox, E.F., Lamb, M.C., Mellentine, S.Q., Tootle, T.L., 2020. Prostaglandins regulate invasive, collective border cell migration. Mol Biol Cell 31, 1584–1594.

Friedl, P., Gilmour, D., 2009. Collective cell migration in morphogenesis, regeneration and cancer. Nat Rev Mol Cell Biol 10, 445–457.

Fulga, T.A., Rorth, P., 2002. Invasive cell migration is initiated by guided growth of long cellular extensions. Nat Cell Biol 4, 715–719.

Gasparski, A.N., Ozarkar, S., Beningo, K.A., 2017. Transient mechanical strain promotes the maturation of invadopodia and enhances cancer cell invasion in vitro. J Cell Sci 130, 1965–1978.

Groen, C.M., Jayo, A., Parsons, M., Tootle, T.L., 2015. Prostaglandins regulate nuclear localization of Fascin and its function in nucleolar architecture. Mol Biol Cell 26, 1901–1917.

Grothey, A., Hashizume, R., Ji, H., Tubb, B.E., Patrick, C.W., Jr., Yu, D., Mooney, E.E., McCrea, P.D., 2000. C-erbB-2/ HER-2 upregulates fascin, an actin-bundling protein associated with cell motility, in human breast cancer cell lines. Oncogene 19, 4864–4875.

Hashimoto, Y., Kim, D.J., Adams, J.C., 2011. The roles of fascins in health and disease. J Pathol 224, 289–300.

Hashimoto, Y., Parsons, M., Adams, J.C., 2007. Dual actin-bundling and protein kinase C-binding activities of fascin regulate carcinoma cell migration downstream of Rac and contribute to metastasis. Mol Biol Cell 18, 4591–4602.

He, L., Wang, X., Tang, H.L., Montell, D.J., 2010. Tissue elongation requires oscillating contractions of a basal actomyosin network. Nat Cell Biol 12, 1133–1142.

Heinrich, H., 1882. Ueber die Berührung fester elastischer Körper. Journal für die reine und angewandte Mathematik 1882, 156–171.

Jayo, A., Malboubi, M., Antoku, S., Chang, W., Ortiz-Zapater, E., Groen, C., Pfisterer, K., Tootle, T., Charras, G., Gundersen, G.G., Parsons, M., 2016. Fascin Regulates Nuclear Movement and Deformation in Migrating Cells. Dev Cell 38, 371–383.

Jayo, A., Parsons, M., 2010. Fascin: a key regulator of cytoskeletal dynamics. Int J Biochem Cell Biol 42, 1614–1617.

Kreplak, L., 2016. Introduction to Atomic Force Microscopy (AFM) in Biology. Curr Protoc Protein Sci 85, 17 17 11–17 17 21.

Kruger, T.M., Bell, K.J., Lansakara, T.I., Tivanski, A.V., Doorn, J.A., Stevens, L.L., 2019. Reduced Extracellular Matrix Stiffness Prompts SH-SY5Y Cell Softening and Actin Turnover To Selectively Increase Abeta(1-42) Endocytosis. ACS Chem Neurosci 10, 1284–1293.

Kruger, T.M., Bell, K.J., Lansakara, T.I., Tivanski, A.V., Doorn, J.A., Stevens, L.L., 2020. A Soft Mechanical Phenotype of SH-SY5Y Neuroblastoma and Primary Human Neurons Is Resilient to Oligomeric Abeta(1-42) Injury. Acs Chem Neurosci 11, 840–850.

Lamb, M.C., Anliker, K.K., Tootle, T.L., 2020. Fascin regulates protrusions and delamination to mediate invasive, collective cell migration in vivo. Dev Dyn 249, 961–982.

Lamb, M.C., Tootle, T.L., 2020. Fascin in Cell Migration: More Than an Actin Bundling Protein. Biology (Basel) 9.

Li, A., Dawson, J.C., Forero-Vargas, M., Spence, H.J., Yu, X., Konig, I., Anderson, K., Machesky, L.M., 2010. The actin-bundling protein fascin stabilizes actin in invadopodia and potentiates protrusive invasion. Curr Biol 20, 339–345.

Li, A., Morton, J.P., Ma, Y., Karim, S.A., Zhou, Y., Faller, W.J., Woodham, E.F., Morris, H.T., Stevenson, R.P., Juin, A., Jamieson, N.B., MacKay, C.J., Carter, C.R., Leung, H.Y., Yamashiro, S., Blyth, K., Sansom, O.J., Machesky, L.M., 2014. Fascin is regulated by slug, promotes progression of pancreatic cancer in mice, and is associated with patient outcomes. Gastroenterology 146, 1386–1396 e1381-1317.

Lo, C.M., Wang, H.B., Dembo, M., Wang, Y.L., 2000. Cell movement is guided by the rigidity of the substrate. Biophys J 79, 144–152.

Ma, Y., Machesky, L.M., 2015. Fascin1 in carcinomas: Its regulation and prognostic value. Int J Cancer 137, 2534–2544.

Majumder, P., Aranjuez, G., Amick, J., McDonald, J.A., 2012. Par-1 controls myosin-II activity through myosin phosphatase to regulate border cell migration. Curr Biol 22, 363–372.

Martinez, J.S., Schlenoff, J.B., Keller, T.C., 3rd, 2016. Collective epithelial cell sheet adhesion and migration on polyelectrolyte multilayers with uniform and gradients of compliance. Exp Cell Res 346, 17–29.

McGowan, S.E., Lansakara, T.I., McCoy, D.M., Zhu, L., Tivanski, A.V., 2020. Platelet-derived Growth Factor-alpha and Neuropilin-1 Mediate Lung Fibroblast Response to Rigid Collagen Fibers. Am J Respir Cell Mol Biol 62, 454–465.

Mierke, C.T., 2020. Mechanical Cues Affect Migration and Invasion of Cells From Three Different Directions. Front Cell Dev Biol 8, 583226.

Minn, A.J., Gupta, G.P., Siegel, P.M., Bos, P.D., Shu, W., Giri, D.D., Viale, A., Olshen, A.B., Gerald, W.L., Massague, J., 2005. Genes that mediate breast cancer metastasis to lung. Nature 436, 518–524.

Mishra, A.K., Mondo, J.A., Campanale, J.P., Montell, D.J., 2019. Coordination of protrusion dynamics within and between collectively migrating border cells by myosin II. Mol Biol Cell 30, 2490–2502.

Mohan, K., Luo, T., Robinson, D.N., Iglesias, P.A., 2015. Cell shape regulation through mechanosensory feedback control. J R Soc Interface 12, 20150512.

Montell, D.J., 2003. Border-cell migration: the race is on. Nat Rev Mol Cell Biol 4, 13–24.

Montell, D.J., Yoon, W.H., Starz-Gaiano, M., 2012. Group choreography: mechanisms orchestrating the collective movement of border cells. Nat Rev Mol Cell Biol 13, 631–645.

Nieto, M.A., Cano, A., 2012. The epithelial-mesenchymal transition under control: global programs to regulate epithelial plasticity. Semin Cancer Biol 22, 361–368.

Oakes, P.W., 2018. Balancing forces in migration. Curr Opin Cell Biol 54, 43–49.

Ono, S., Yamakita, Y., Yamashiro, S., Matsudaira, P.T., Gnarra, J.R., Obinata, T., Matsumura, F., 1997. Identification of an actin binding region and a protein kinase C phosphorylation site on human fascin. J Biol Chem 272, 2527–2533.

Parekh, A., Weaver, A.M., 2016. Regulation of invadopodia by mechanical signaling. Exp Cell Res 343, 89–95.

Patel, N.H., Snow, P.M., Goodman, C.S., 1987. Characterization and cloning of fasciclin III: a glycoprotein expressed on a subset of neurons and axon pathways in Drosophila. Cell 48, 975–988.

Platt, J.L., Michael, A.F., 1983. Retardation of fading and enhancement of intensity of immunofluorescence by p-phenylenediamine. J Histochem Cytochem 31, 840–842.

Ren, Y., Zhang, Y., Liu, J., Liu, P., Yang, J., Guo, D., Tang, A., Tao, J., 2021. Matrix hardness regulates the cancer cell malignant progression through cytoskeletal network. Biochem Biophys Res Commun 541, 95–101.

Roubinet, C., Tsankova, A., Pham, T.T., Monnard, A., Caussinus, E., Affolter, M., Cabernard, C., 2017. Spatio-temporally separated cortical flows and spindle geometry establish physical asymmetry in fly neural stem cells. Nat Commun 8, 1383.

Shellard, A., Mayor, R., 2021. Durotaxis: The Hard Path from In Vitro to In Vivo. Dev Cell 56, 227–239.

Spradling, A., 1993. Developmental genetics of oogenesis. Cold Spring Harbor Laboratory Press, The development of Drosophila melanogastor, pp. 1–70.

Stuelten, C.H., Parent, C.A., Montell, D.J., 2018. Cell motility in cancer invasion and metastasis: insights from simple model organisms. Nat Rev Cancer 18, 296–312.

Sunyer, R., Conte, V., Escribano, J., Elosegui-Artola, A., Labernadie, A., Valon, L., Navajas, D., Garcia-Aznar, J.M., Munoz, J.J., Roca-Cusachs, P., Trepat, X., 2016. Collective cell durotaxis emerges from long-range intercellular force transmission. Science 353, 1157–1161.

Sunyer, R., Trepat, X., 2020. Durotaxis. Curr Biol 30, R383–R387.

Telley, I.A., Gaspar, I., Ephrussi, A., Surrey, T., 2012. Aster migration determines the length scale of nuclear separation in the Drosophila syncytial embryo. J Cell Biol 197, 887–895.

Tse, J.M., Cheng, G., Tyrrell, J.A., Wilcox-Adelman, S.A., Boucher, Y., Jain, R.K., Munn, L.L., 2012. Mechanical compression drives cancer cells toward invasive phenotype. Proc Natl Acad Sci U S A 109, 911–916.

van Helvert, S., Friedl, P., 2016. Strain Stiffening of Fibrillar Collagen during Individual and Collective Cell Migration Identified by AFM Nanoindentation. ACS Appl Mater Interfaces 8, 21946–21955.

Vicente-Manzanares, M., Ma, X., Adelstein, R.S., Horwitz, A.R., 2009. Non-muscle myosin II takes centre stage in cell adhesion and migration. Nat Rev Mol Cell Biol 10, 778–790.

Villari, G., Jayo, A., Zanet, J., Fitch, B., Serrels, B., Frame, M., Stramer, B.M., Goult, B.T., Parsons, M., 2015. A direct interaction between fascin and microtubules contributes to adhesion dynamics and cell migration. J Cell Sci 128, 4601–4614.

Yamakita, Y., Ono, S., Matsumura, F., Yamashiro, S., 1996. Phosphorylation of human fascin inhibits its actin binding and bundling activities. J Biol Chem 271, 12632–12638.

Zaccai, M., Lipshitz, H.D., 1996. Differential distributions of two adducin-like protein isoforms in the Drosophila ovary and early embryo. Zygote 4, 159–166.

Zanet, J., Jayo, A., Plaza, S., Millard, T., Parsons, M., Stramer, B., 2012. Fascin promotes filopodia formation independent of its role in actin bundling. J Cell Biol 197, 477–486.

Zanet, J., Stramer, B., Millard, T., Martin, P., Payre, F., Plaza, S., 2009. Fascin is required for blood cell migration during Drosophila embryogenesis. Development 136, 2557–2565.

